# Morphogenesis of the muscular ventricular septum of the mouse heart is driven by retinoic acid signalling to myocardium

**DOI:** 10.64898/2026.08.12.744354

**Authors:** Tobias H. Bønnelykke, Célia Coulon, Rachel Sturny, Marie Couderc, Claudio Cortés, Célia Rousset, Diptarka Saha, Damien Marchese, Christopher De Bono, Lucile Miquerol, Gonzalo del Monte-Nieto, Stéphane Zaffran, Robert G. Kelly

## Abstract

The mammalian heart is divided into four chambers by septa that isolate systemic and pulmonary circulation and are hotspots of congenital heart defects (CHD). The muscular ventricular septum develops between left and right ventricular cardiomyocytes derived from the first and second heart fields. Despite its clinical importance, mechanisms underlying development of the ventricular septum are poorly understood. Here we show that myocardial reception of retinoic acid (RA) signalling regulates formation of the compact septal core. Activation of a dominant negative RA receptor in second heart field-derived myocardium during septal morphogenesis results in a deep interventricular cleft and bifid cardiac apex. This phenotype is preceded by ectopic trabecular contributions to a RA-independent septal primordium. Molecular analysis implicates defective cardiomyocyte maturation and impaired RAC1 activation in mutant hearts. These results support an infolding and RA-dependent fusion model of septal morphogenesis, providing new insights into ventricular development and the origins of CHD.

## Introduction

The left and right sides of the mammalian heart are divided by cardiac septa that ensure separation of oxygen rich and poor blood and are hotspots of congenital heart defects. The interventricular septum (IVS) is comprised of a large muscular part and a membranous component at the junction with the outflow tract and atrioventricular cushions^1^. Ventricular septal anomalies account for over 35% of congenital heart defects and arise during morphogenesis of the IVS that takes place between the 5th and 8th week of human development and embryonic days (E) 10.5 and 14.5 in the mouse^2, 3^.

The muscular IVS forms at the interface between cardiomyocytes derived from the first and second heart field^4, 5^. Molecular and lineage analyses have demonstrated that the muscular septum contains cardiomyocytes of left and right ventricular identity separated by a border that acts as a clonal compartment boundary within the developing heart^4, 6–8^. Genetic studies have shown that the site of formation of the ventricular septum is prepatterned in the early heart and that a sharp gradient of *Tbx5* expression is required for septum morphogenesis^9, 10^. Embryonic ventricular myocardium is composed of sub-endocardial trabeculae, finger-like projections of myocardium, and a compact myocardial layer that forms the mass of the definitive ventricular muscle and constitutes the central core of the muscular IVS^11, 12^. Based on detailed human and comparative anatomical studies, experimental embryology in avians and analyses of mouse mutants with congenital heart defects, several different models of IVS formation have been proposed. These include outgrowth from the cardiac apex towards the base, coalescence of central trabecular myocardium, or folding, apposition and fusion of compact layer myocardium during ventricular chamber morphogenesis^4, 13–20^. Yet to date, despite the clinical importance of ventricular septation, the mechanisms driving formation of the muscular IVS are poorly understood.

The Vitamin A derivative retinoic acid (RA) plays diverse essential roles in patterning and differentiation during embryonic development through transcriptional activation at RA-response elements mediated by interaction with nuclear receptors of the RAR and RXR families^21^. RA signalling has long been recognized to play important roles at multiple stages of heart development^22^, including patterning and specifying early cardiac progenitor cells, delimiting the posterior boundary of the second heart field, and regulating progenitor cell deployment to the arterial and venous poles of the heart^22–24^. Vitamin A deficiency also results in a range of later cardiac defects including ventricular wall hypoplasia^25^. Similar defects, including hypoplasia of the compact ventricular wall and a trabeculated septum, are seen in mouse embryos lacking RXRA or carrying combinations of RAR and RXR mutations^26–29^. RA is produced in the overlying outer layer of the heart, the epicardium, where retinaldehyde dehydrogenases necessary for RA synthesis are expressed during chamber morphogenesis^30^. However, a series of genetic experiments have suggested that epicardially synthesized RA does not signal directly to the adjacent compact myocardial layer. Instead, autocrine epicardial RA signalling and extra-cardiac liver sources of RA-dependent Erythropoetin regulate the expression of mitogens, including FGFs and IGF2, in the epicardium that in turn signal to the adjacent myocardium to promote compact myocardial proliferation and regulate the timing of myocardial differentiation^31–36^. Moreover, RA-dependent epicardial signalling and epicardial derived smooth muscle and fibroblast contributions to the ventricle play major roles in establishing the coronary vasculature^37, 38^. Different genetic systems using reporter genes driven by RA-response elements (RARE) have been developed to identify the cellular targets of RA^30, 35, 39, 40^. While these studies have pointed to the importance of the epicardial response to RA, analysis of an inducible *RARE-CreERT2* transgene has revealed that compact layer cardiomyocytes respond directly to RA signalling after E10.5, during the time of compact wall growth and septal morphogenesis^41^. The role of direct RA signalling to the developing myocardium, however, remains unknown.

Using genetic and pharmacological approaches we show that RA signal reception is required in myocardial cells contributing to the compact core of the septum for fusion of apposed medial left and right ventricular chamber walls, revealing a previously unrecognised role for RA signalling directly to ventricular myocardium in formation of the muscular IVS. These results support an infolding and RA-dependent fusion model of ventricular septal morphogenesis. Failure of this process results in formation of a bifid ventricular apex and deep interventricular cleft. In addition, RA signalling to the myocardium is required for cardiomyocyte maturation and normal coronary artery development. Our results provide new insights into ventricular septation and the complexity of RA signalling during cardiac development.

## Results

### Expression of a dominant negative retinoic acid receptor in the second heart field lineage leads to failure of muscular interventricular septum morphogenesis

In order to investigate the cell autonomous requirement for retinoic acid (RA) signal reception in the second heart field (SHF) and SHF-derived parts of the developing heart, mice carrying *RARα403*, a conditional dominant negative RA receptor, were crossed with *Mef2c-AHF-Cre* mice (Fig. 1a). The *Mef2c-AHF-Cre* genetic lineage includes SHF progenitor cells and their myocardial derivatives in the outflow tract, right ventricle and interventricular septal (IVS) region (Figs. 1b and S1a; Video S1)^42^. *R26^RARα403^* encodes a truncated human RA receptor lacking the C-terminal ligand-dependent transcriptional activation domain, inserted at the ubiquitous *Rosa26* locus and preceded by a floxed stop codon, allowing Cre-dependent cell autonomous reduction of RAR-mediated RA signalling^43^. Fetal hearts were analysed at embryonic day (E) 14.5. Wholemount analysis revealed a striking defect in ventricular shape in *Mef2c-AHF-Cre^Tg+^;R26^RARα403^* hearts. The right and left ventricular chambers displayed a rounded ventricular morphology associated with a failure of apical convergence (Fig. 1c). An interventricular cleft running from the ventral to the dorsal surface of the heart was observed in all *Mef2c-AHF-Cre^Tg+^;R26^RARα403^* hearts analysed but not in hearts carrying either *Mef2c-AHF-Cre^Tg+^* or *R26^RARα403^* alone. Histological analysis revealed that while the IVS is fully formed at this stage in control hearts, *Mef2c-AHF-Cre^Tg+^;R26^RARα403^* hearts lack a normal muscular IVS and instead display a deep cleft between juxtaposed right and left chamber walls, below a residual muscular septum (Fig. 1d). The lack of normal apposition of the right and left chambers results in a bifid ventricular morphology with two well separated ventricular apexes. This reveals a previously unappreciated requirement for RA signal reception in the SHF lineage for IVS morphogenesis. In addition, while outflow tract division to form the ascending aorta and pulmonary trunk is complete at E14.5 in control hearts, this process fails in *Mef2c-AHF-Cre^Tg+^;R26^RARα403^* hearts, resulting in a common arterial trunk (Fig. 1c,d)^44^.

**Figure 1.**
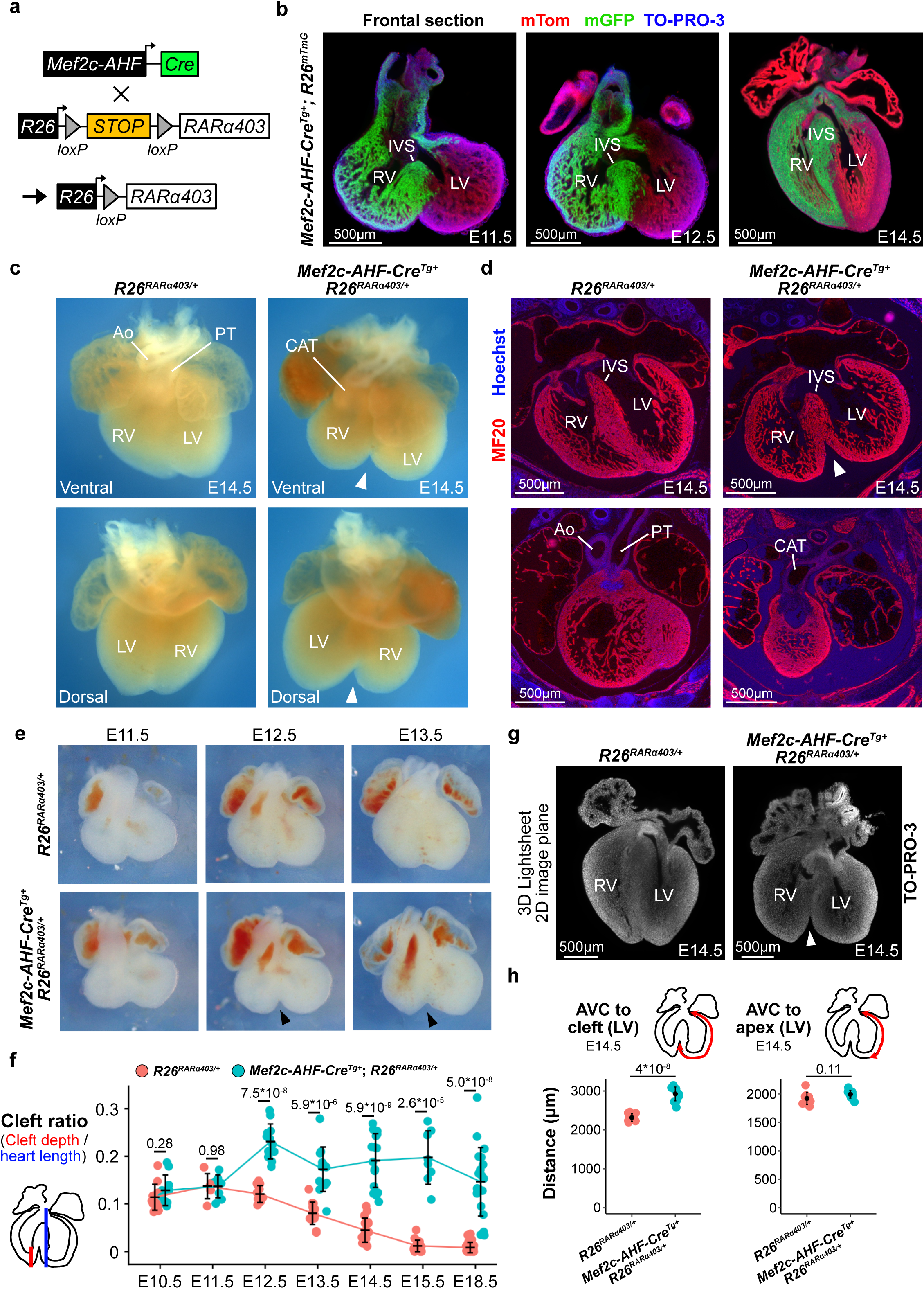
Expression of a dominant negative retinoic acid receptor in the *Mef2c-AHF-Cre* genetic lineage results in a bifid ventricular morphology. **(a)** Scheme of the genetic cross leading to *Mef2c-AHF-Cre* dependent activation of the dominant-negative RA receptor gene *RARα403*. **(b)** *Mef2c-AHF-Cre* lineage contributions (green) in frontal sections of wholemount fluorescent lightsheet images of cleared *Mef2c-AHF-Cre^Tg+^*;*R26^mTmG^* hearts at E11.5, E12.5 and E14.5. **(c)** Ventral and dorsal brightfield images of E14.5 control *R26^RARα403^* and mutant *Mef2c-AHF-Cre^Tg+^*;*R26^RARα403^* hearts. Mutant hearts have a common arterial trunk (CAT) and a deep interventricular cleft (arrowheads). **(d)** Frontal sections of E14.5 *R26^RARα403^* and *Mef2c-AHF-Cre^Tg+^*;*R26^RARα403^* hearts labelled with MF20 (red), highlighting the interventricular cleft (arrowhead) and the CAT in mutant hearts **(e)** Ventral brightfield images of *R26^RARα403^* and *Mef2c-AHF-Cre^Tg+^*;*R26^RARα403^* hearts from E11.5 to E13.5. Control hearts initially have an interventricular cleft, that is resolved during fetal heart development. Black arrowheads indicate the abnormal interventricular clefts in mutant hearts. **(f)** Quantification of bifid severity in *R26^RARα403^* control and *Mef2c-AHF-Cre^Tg+^*;*R26^RARα403^* mutant hearts from stages E10.5 to E18.5. Cleft ratio diverges between control and mutants from E12.5. Means, standard deviations and P-values (Welch’s t-test) are shown. N [Control; Mutant]: E10.5 [12;10]; E11.5 [6;8]; E12.5 [13;12]; E13.5 [14;13]; E14.5 [16;16]; E15.5 [15;8]; E18.5 [25;20]. **(g)** Frontal sections of wholemount *R26^RARα403^*and *Mef2c-AHF-Cre^Tg+^*;*R26^RARα403^* hearts at E14.5; nuclei are fluorescently labelled using TO-PRO-3. **(h)** Measurement of the left ventricular perimeter between the AVC and cleft (left) and AVC and left ventricular apex (right) in E14.5 *R26^RARα403^* and *Mef2c-AHF-Cre^Tg+^*;*R26^RARα403^* hearts, showing means, standard deviations and P-values (Welch’s t-test). N [Control; Mutant]: left panel [10;11], right panel [9;10]. Ao, Aorta; AVC, atrioventricular canal; IVS, interventricular septum; LV, left ventricle; PT, pulmonary trunk; RV, right ventricle.

The emergence of the bifid ventricular phenotype was investigated between E10.5 and E18.5 (Figs. 1e,f; S1b-d). In control hearts, an interventricular apical cleft present at E10.5 gradually disappears after E13.5. *Mef2c-AHF-Cre^Tg+^;R26^RARα403^* hearts at E10.5 and E11.5 are similar to those of control littermates, however from E12.5 the interventricular cleft increases in size and fails to be resolved during subsequent development. To quantify the interventricular cleft phenotype, we calculated the ratio between cleft size and ventricular base to apex length (Fig. 1f). Quantification confirmed a highly significant divergence in this ratio between control and mutant hearts from E12.5. In addition to an apical interventricular cleft, late fetal stage (E18.5) *Mef2c-AHF-Cre^Tg+^;R26^RARα403^* hearts displayed a degree of hypoplasia of the right ventricular wall (observed in 8/20 hearts) while 2/20 *Mef2c-AHF-Cre^Tg+^;R26^RARα403^* hearts appeared enlarged (Fig. S1c). No *Mef2c-AHF-Cre^Tg+^;R26^RARα403^*mice were recovered after birth, likely due to failure to separate systemic from pulmonary circulation as a result of the common arterial trunk. To complement our 2D brightfield image-based quantification of cleft length, we collected hearts at E14.5 and performed cleared 3D lightsheet imaging (Fig. S1e; Video S2). From these images, we obtained precisely matching 2D planes for control and mutant hearts (Fig. 1g). Quantitative analysis confirmed a significant increase in cleft length in *Mef2c-AHF-Cre^Tg+^;R26^RARα403^* hearts with no difference in heart length (Fig. S1f). Further quantification revealed that while the distance from the AVC to the left ventricular apex is similar in control and *Mef2c-AHF-Cre^Tg+^;R26^RARα403^* hearts, the distance from the left ventricular apex to the top of the cleft is increased (Fig. 1h). Similarly, the distance between the apex of the right and left ventricular chambers is increased in mutant hearts at E14.5 (Fig. S1f). These results are consistent with a failure of apposed medial right and left ventricular walls to adhere in the apical muscular septum. To rule out that the smaller septum was not due to defects in proliferation, we investigated expression of the proliferation marker Ki67 in E12.5 hearts (Fig. S2a). Septal area, cell number, cell density and proliferation index were quantified separately for myocardial and endocardial cells. While the IVS of *Mef2c-AHF-Cre^Tg+^;R26^RARα403^* hearts is smaller with less cells, the ratio of Ki67+ cells remained the same as in *R26^RARα403^* hearts without *Cre*, indicating no changes in proliferation in either septal myocardium or endocardium (Fig. S2b). Immunofluorescence detection of cleaved Caspase-3 revealed similar low levels of labelling in the septal region of control and *Mef2c-AHF-Cre^Tg+^;R26^RARα403^* hearts at E13.5, suggesting that the bifid cleft does not result from altered levels of apoptotic cell death (Fig. S2c).

### Reduced RA signalling does not affect early patterning of the interventricular region but directly impacts the later process of septum formation

We next confirmed that RA signalling was reduced in the *Mef2c-AHF-Cre* genetic lineage in *Mef2c-AHF-Cre^Tg+^;R26^RARα403^* hearts. Analysis of β-galactosidase expression driven by a *RARE-lacZ* transgene ^39^ revealed reduced transgene activity in the right ventricle of *Mef2c-AHF-Cre^Tg+^;R26^RARα403^* compared to control *R26^RARα403^* hearts at E12.5 (Fig. S3a). Histological sections showed that *RARE-lacZ* is expressed in both epicardium and outer layer compact myocardium at E12.5 (Fig. S3b), in agreement with previous results ^30^. In *Mef2c-AHF-Cre^Tg+^;R26^RARα403^* hearts, right ventricular myocardial, but not epicardial, β-galactosidase labelling was reduced (Fig. S3b), consistent with the myocardial-specific contribution of the *Mef2c-AHF-Cre* lineage (Fig.1a; ^42^). The *Mef2c-AHF-Cre* lineage overlaps with *Tbx5* expressing cells in the left side of the future interventricular region^7, 8, 42^. To explore if RA signal reception is required for molecular patterning of the interventricular region, we generated lineage traced control and *Mef2c-AHF-Cre^Tg+^;R26^RARα403^* hearts using the *Z/EG* conditional reporter gene^45^. The extent of *Mef2c-AHF-Cre* lineage contributions between control and mutant hearts was similar at E10.5 (Figs. 2a and S4a). Quantification of sections of 3D imaged hearts at E11.5 revealed an indistinguishable extent of GFP labelling in the septal region in *Mef2c-AHF-Cre^Tg+^;R26^RARα403^;Z/EG* and *Mef2c-AHF-Cre^Tg+^;Z/EG* hearts (Fig. S4b-d). At E14.5, when the cleft phenotype is evident, myocardium on the left ventricular side of the cleft is negative for the *Mef2c-AHF-Cre* genetic lineage, while the contribution to the residual septum resembles that of control hearts (Fig. 2b). RNAscope hybridization revealed similar patterns of *Tbx5* transcript distribution in *Mef2c-AHF-Cre^Tg+^;R26^RARα403^* and control hearts at E13.5 (Fig. S4e). Together, these results suggests that molecular patterning of the interventricular region is established normally in *Mef2c-AHF-Cre^Tg+^;R26^RARα403^* hearts. In further support of normal patterning, the expression domain of *Tbx18* on the left side of the residual muscular septum is retained in *Mef2c-AHF-Cre^Tg+^;R26^RARα403^* hearts (Fig. S4f)^4^. However, *Tbx18* transcripts are also observed in the left ventricular wall close to the residual septum of *Mef2c-AHF-Cre^Tg+^;R26^RARα403^* hearts at E13.5, consistent with failure of normal apposition of left and right septal proximal myocardium (Fig. S4f, arrowhead).

**Figure 2.**
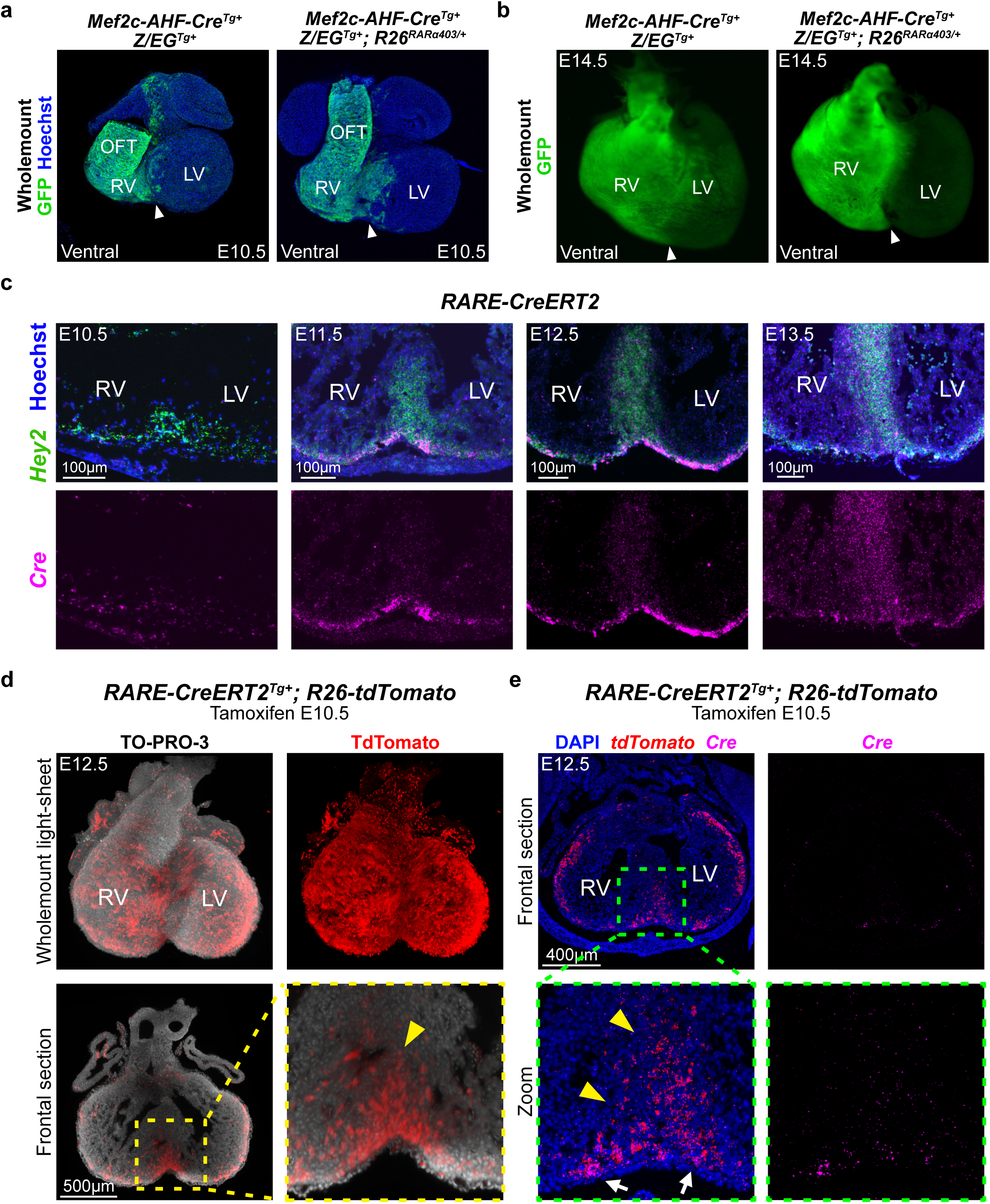
Retinoic acid signal reception in the *Mef2c-AHF-Cre* genetic lineage does not affect early patterning of the interventricular region and is observed during subsequent septal morphogenesis. **(a)** Maximum intensity projection (MIP) of wholemount fluorescent labelled *Mef2c-AHF-Cre^Tg+^;Z/EG^Tg^*^+^ and *Mef2c-AHF-Cre^Tg+^;Z/EG^Tg+^;R26^RARα403^* hearts at E10.5 in ventral views showing the contribution of the *Mef2c-AHF-Cre* genetic lineage. **(b)** MIP of wholemount fluorescent labelled *Mef2c-AHF-Cre^Tg+^;Z/EG^Tg^*^+^ and *Mef2c-AHF-Cre^Tg+^;Z/EG^Tg+^;R26^RARα403^* hearts at E14.5. Similar contributions are observed in control and mutant hearts. **(c)** RNAscope labelling of *Hey2* (green) and *Cre* (magenta) transcripts in sectioned *RARE-CreERT2^Tg+^* hearts from E10.5 to E13.5 focused on the septal region. Strong *Cre* expression is observed in *Hey2* expressing cells in the outer compact ventricular myocardial layer from E11.5. **(d)** Wholemount and frontal sections of *RARE-CreERT2^Tg+^*;*R26^tdTomato^* hearts at E12.5 after tamoxifen injection at E10.5. In contrast to *Cre* transcripts, tdTomato^+^ cells are observed in the ventricular septum (arrowhead). N: 2 litters, 6 hearts. **(e)** Colabelling of *tdTomato* (red) and *Cre* (magenta) by RNAscope in sectioned E12.5 *RARE-CreERT2^Tg+^* hearts after tamoxifen administration at E10.5. *Cre* expression is observed in the outer compact ventricular myocardial layer (arrow), while *tdtomato* expression is observed within the septum (arrowhead). LV, left ventricle; OFT, outflow tract; RV, right ventricle

We next investigated the temporal requirement for RA signal reception in muscular ventricular septal morphogenesis. Prior experiments in which RA signalling was pharmacologically inhibited at E8.75 and E9.25 did not result in bifid apex hearts^24^. It has recently been shown using a *RARE-CreERT2* transgene that the compact myocardium responds to RA signalling from E10.5^41^. To explore the kinetics of RA signal response in the myocardium, we performed RNAscope *in situ* hybridization for *Cre* transcripts in *RARE-CreERT2^Tg+^* mice^41^ from E10.5 to E13.5 along with a marker of compact myocardium, *Hey2* (Figs. 2c and S5a). Consistent with the findings of Da Silva *et al*. ^41^, we observed that C*re* transcripts accumulate in compact ventricular myocardium at E11.5 and E12.5, while expression in the compact core of the IVS is observed only after E13.5. These results suggest that the compact layer of ventricular myocardium is RA responsive at the time of ventricular septum formation. The epicardium expands over the developing ventricles from E10.5 and expresses the aldehyde dehydrogenase gene *Aldh1a2*, and is thus a source of RA adjacent to compact ventricular myocardium^30^. Epicardial expression of both *Aldh1a1* and *Aldh1a2* was confirmed using published single cell RNA-seq data of mouse hearts from E10.5 to E14.5^46^ (Fig. S5b). Moreover, *Hey2^+^* compact ventricular cardiomyocytes express the genes encoding the retinoic acid receptors *Rara* and *Rarb* and the retinoid X receptors *Rxra* and *Rarb* (Fig. S5c). We next investigated the fate of RA-responding cells in compact myocardium in E12.5 *RARE-CreERT2^Tg+^*;*R26^tdTomato^* hearts following tamoxifen injection at E10.5 (Fig. 2d; Video S3). Lightsheet imaging revealed TdTomato labelling throughout the ventricular compact myocardium, including extended labelling in the compact core of the lower part of the ventricular septum (arrowhead in Fig. 2d). Dual analysis of *Cre* and *Tomato* transcript distribution in *RARE-CreERT2^Tg+^*;*R26^tdTomato^* hearts confirmed accumulation of *Tomato* transcripts in the septal core at E12.5, while *Cre* expression is restricted to subepicardial myocardium (Fig. 2e). These results suggest that RA-responding sub-epicardial compact myocardium contributes to the muscular septal core.

Having shown that compact ventricular cardiomyocytes respond to RA signalling and contribute to the IVS, we next asked whether pharmacological inhibition of RA signalling during septal morphogenesis would phenocopy the *Mef2c-AHF-Cre^Tg+^;R26^RARα403^* ventricular septal phenotype. The pan-RAR inverse agonist BMS493 was injected into pregnant wildtype CD1 female mice for 3 consecutive days between E11.5 and E13.5 and the bifid cleft ratio quantified in dissected hearts at E14.5 (Fig. 3a-c). This treatment induced a bifid ventricular phenotype, although with incomplete penetrance compared to the genetic reduction of RA signal reception (Fig. 1). We also performed injections for 2 consecutive days, which led to significant, albeit milder, increases in cleft ratio compared to PBS-injected controls (Fig. S6a,b). These results confirm that RA signal reception is required for septal morphogenesis during the process of ventricular septum formation itself rather than at an earlier patterning step. Together these data support an infolding model for formation of the muscular IVS during chamber ballooning morphogenesis, whereby ventricular compact myocardial cells respond to RA signalling and integrate into the septal core.

**Figure 3.**
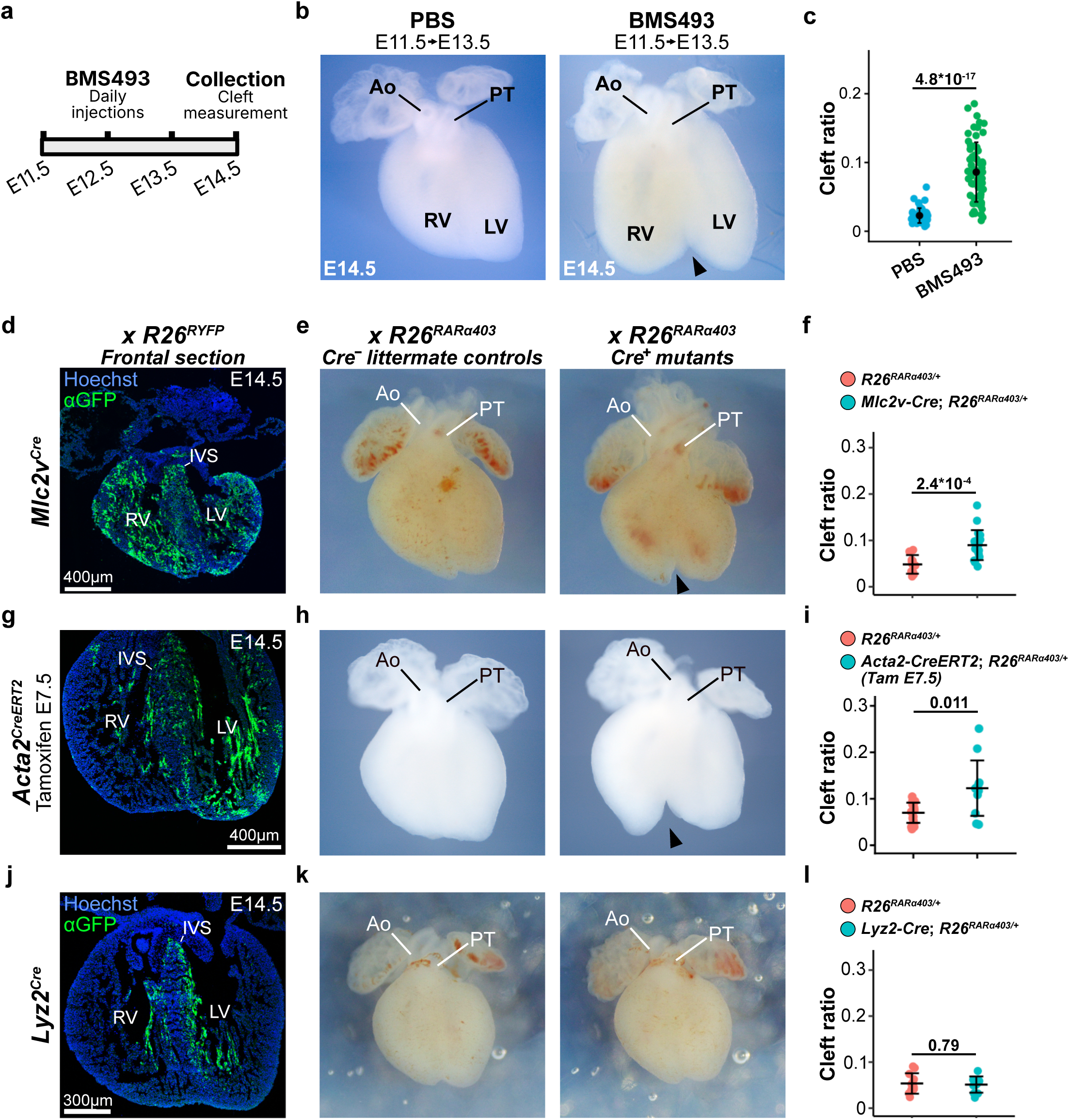
Temporal and spatial refinement of the requirement for retinoic acid reception for ventricular septal morphogenesis. **(a)** Scheme showing the experimental procedure of inverse pan-RAR agonist BMS493 injection on 3 consecutive days from E11.5 to E13.5 followed by collection of hearts and scoring of cleft ratio at E14.5. **(b)** Brightfield images of PBS treated and BMS493 treated (E11.5+E12.5+E13.5) hearts at 14.5; the black arrowhead indicates a bifid cleft. **(c)** Cleft ratio measurements of E14.5 PBS treated- and BMS493 treated hearts showing means, standard deviations and P-value (Welch’s t-test). N: [PBS = 50; BMS493 = 62. **(d-f)** *Mlc2v^Cre^*;*R26^RYFP^* lineage contributions at E14.5 in a GFP-stained section (d), littermate control and *Mlc2v^Cre^;RARa403* E14.5 hearts (e) and quantification of cleft ratio (f); **(g-i)** *Acta2^CreERT2^*;*R26^RYFP^* (tamoxifen E7.5) lineage contributions at E14.5 in a GFP-stained section (g), littermate control and *Acta2^CreERT2^;RARa403* (tamoxifen E7.5) E14.5 hearts (h) and quantification of cleft ratio (i). (j-l) *Lyz2^Cre^*;*R26^RYFP^* lineage contributions at E14.5 in a GFP-stained section (j), littermate control and *Lyz2^Cre^;RARa403* E14.5 hearts (k) and quantification of cleft ratio (l). Black arrowheads indicate bifid clefts. Means, standard deviations and P-value (Welch’s t-test) are shown. N = [Control; Mutant] : *Mlc2v^Cre^* [10;19], *Acta2^CreERT2^* [17;12], *Lyz2^Cre^* [11;10]. Ao, Aorta; IVS, interventricular septum; LV, left ventricle; RV, right ventricle; PT, pulmonary trunk.

### Retinoic acid signal reception is required in myocardial cells contributing to the ventricular septal core for normal septum morphogenesis

We next addressed the spatial requirement for RA signalling in septum morphogenesis by activating RARα403 with different myocardial *Cre* lines and evaluating lineage contributions and ventricular shape at E14.5. We first used a *Mlc2v^Cre^* transgene that drives Cre recombinase activity in ventricular myocardium (Fig. 3d)^31^. *Mlc2v^Cre^;R26^RARα403^* hearts displayed a bifid ventricular apex at E14.5, albeit with partial penetrance and a less extensive interventricular cleft, likely due to variegated recombination in myocardium (Fig. 3d-f). In contrast to the situation in *Mef2c-AHF-Cre^Tg+^;R26^RARα403^* hearts, outflow tract division occurs normally in *Mlc2v^Cre^;R26^RARα403^* hearts, dissociating these two aspects of the *Mef2c-AHF-Cre^Tg+^;R26^RARα403^* phenotype (Fig. 3e). We then investigated whether RA-response is required specifically in septal myocardium for normal IVS morphogenesis using an *Acta2^CreERT2^* allele^47^. *Acta2*, encoding smooth muscle actin, is expressed in early differentiating cardiomyocytes and induction of recombinase activity at E7.5 results in activation of a conditional YFP reporter gene in the early heart tube and later in septal and left ventricular myocardium at E14.5 (Fig. 3g; ^48^). *Acta2^CreERT2^;R26^RARα403^* hearts displayed a bifid ventricular apex at E14.5, albeit with partial penetrance likely due to Cre mosaicism, and normal outflow tract septation (Fig. 3h,i). *Lyz2*, encoding Lysozyme M, is transiently expressed in the interventricular septal primordium of the embryonic heart^9^. The *Lyz2^Cre^* lineage labels the trabecular component of the IVS with only a minor cellular contribution to the compact septal core (Fig. 3j). Investigation of cardiac structure showed that *Lyz2^Cre^;R26^RARα403^* hearts do not have a bifid ventricular phenotype at E14.5 (Fig. 3k,l). Together with the *Mef2c-AHF-Cre^Tg+^;R26^RARα403^* phenotype, these results identify myocardium contributing to the compact, and not trabecular, region of the IVS as the target of RA signalling during septal morphogenesis. Subsequent experiments focus on *Mef2c-AHF-Cre^Tg+^;R26^RARα403^* hearts which have a fully penetrant bifid ventricular phenotype.

### Expansion of trabecular myocardium in the interventricular septum on downregulation of retinoic acid signal reception

In order to obtain molecular insights into the impact of *RARα403* expression in myocardium, we performed single cell RNA sequencing of *Mef2c-AHF-Cre^Tg+^;R26^RARα403^* and control hearts at E11.5 (Fig. 4). This timepoint, prior to the emergence of the bifid ventricular phenotype, corresponds to the onset of myocardial RA response and ventricular septal morphogenesis.

**Figure 4.**
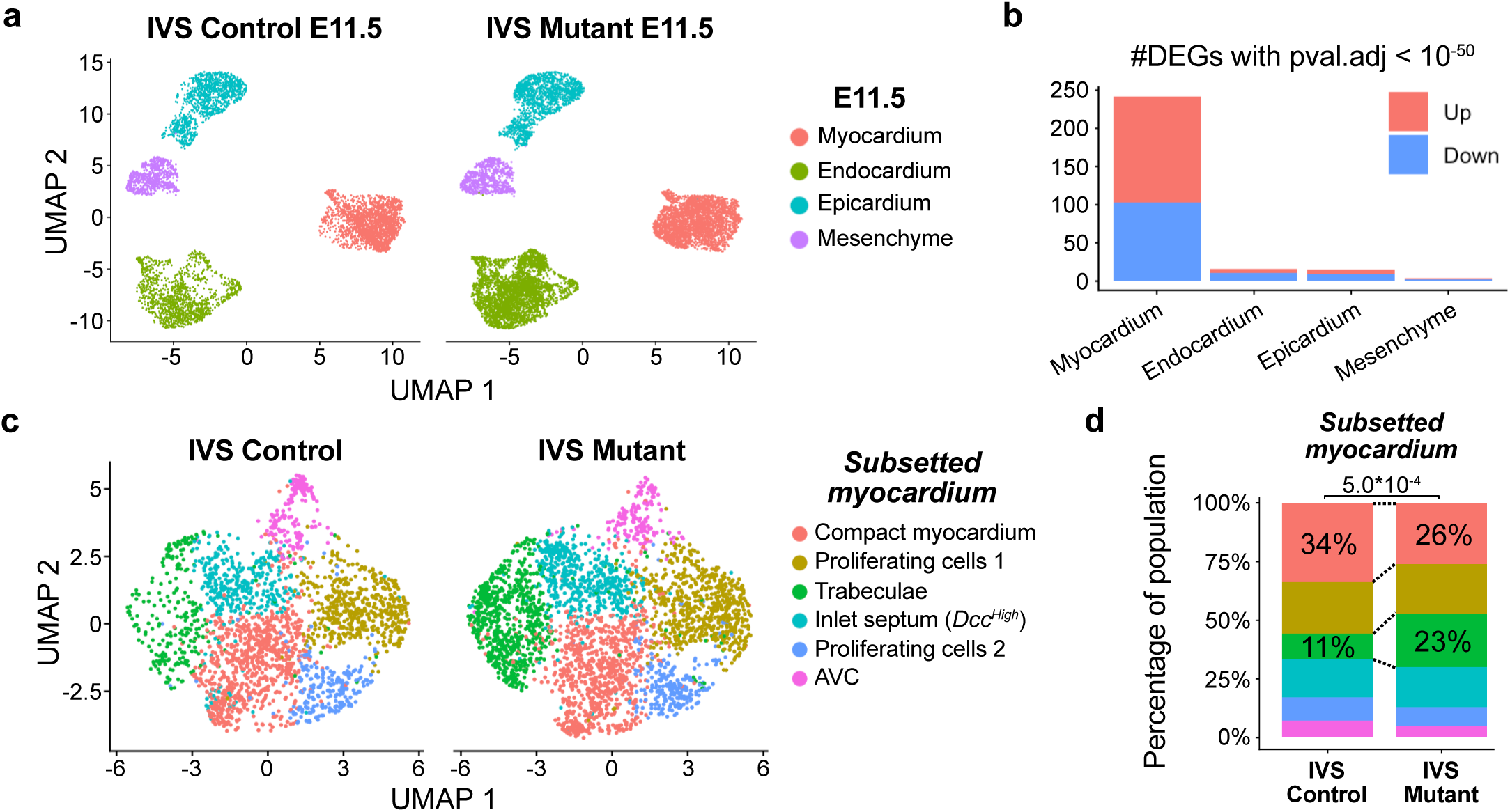
Single cell transcriptomic analysis identifies retinoic acid dependent changes in the myocardium of *Mef2c-AHF-Cre^Tg+^*;*R26^RARα403^* hearts. **(a)** UMAP showing annotated cell clusters from micro-dissected E11.5 IVS regions of *R26^RARα403^*and *Mef2c-AHF-Cre^Tg+^*;*R26^RARα403^* hearts. **(b)** Bar plot showing number of DEGs per clusters with p-value < 10^-50^; colours indicating up- and downregulated DEGS in mutant hearts. **(c)** UMAP showing annotated subclusters of subsetted myocardial cells. **(d)** Stacked bar plots showing the distribution of cells in myocardial subclusters between control and mutant hearts. P-value (simulated Fisher’s test, 2000 iterations) show differences in the cell type distribution between genotypes. Notably, in the mutant IVS the number of compact myocardial cells is decreased and the number of trabecular cells increased. DEG, differentially expressed gene; IVS, interventricular septum; UMAP, Uniform Manifold Approximation and Projection.

Microdissected IVS regions of *Mef2c-AHF-Cre^Tg+^;R26^RARα403^*and control *R26^RARα403^* hearts were processed for scRNA-seq and annotation identified myocardial, endocardial, epicardial and mesenchymal cell clusters (Figs. 4a and S7a-c). Differentially expressed gene (DEG) analysis between control and mutant cells of the distinct clusters revealed that the largest number of significant DEGs was in the myocardium (Figs. 4b and S7d-g; Table S1), consistent with *Mef2c-AHF-Cre* lineage contributions and the distribution of RA-responding cells observed by *Cre* expression in *RARE-CreERT2^Tg+^* hearts.

Subclustering of myocardial cells revealed distinct subclusters corresponding to compact myocardium, trabeculae, atrioventricular region myocardium and proliferating cells (Figs. 4c and S7c). Significant differences were observed between control and mutant samples: the proportion of cells in the compact myocardial cluster was reduced and that in the trabecular cluster increased in *Mef2c-AHF-Cre^Tg+^;R26^RARα403^* hearts (Fig. 4d). Multiple downregulated compact myocardial genes and upregulated trabecular genes were observed in *Mef2c-AHF-Cre^Tg+^;R26^RARα403^* myocardial cells (Fig. S8a). These results were validated in vivo using RNAscope in situ hybridisation with *Hey2* and *Nppa* to label compact and trabecular myocardium respectively (Fig. 5a,b; Video S4). In wildtype hearts, the E10.5 septum primordium extended from E11.5 to form a *Hey2* expressing compact septum. In *Mef2c-AHF-Cre^Tg+^;R26^RARα403^* hearts, the E10.5 septum primordium fails to compact at E11.5, leading to a septal core with ectopic expression of *Nppa* and a more trabeculated appearance (Fig. 5a,b arrowheads). Additional quantification revealed that the thickness of the compact myocardium of the right ventricular free wall is decreased in *Mef2c-AHF-Cre^Tg+^;R26^RARα403^* hearts from E12.5 (Fig. S8b-d), suggesting that myocardial RA reception plays a broader role in growth of the ventricular wall. *Hey2* and *Nppa* expression were analysed by RNAscope hybridisation in sections of bifid ventricular hearts generated by BMS493 exposure between E11.5 and E13.5. No ectopic trabecular gene expression was observed in the septal core of these hearts (Fig. 5c). This result suggests that expanded trabecular myocardium in the septal core and the bifid ventricular cleft are temporally separate phenotypes resulting from decreased RA signalling.

**Figure 5.**
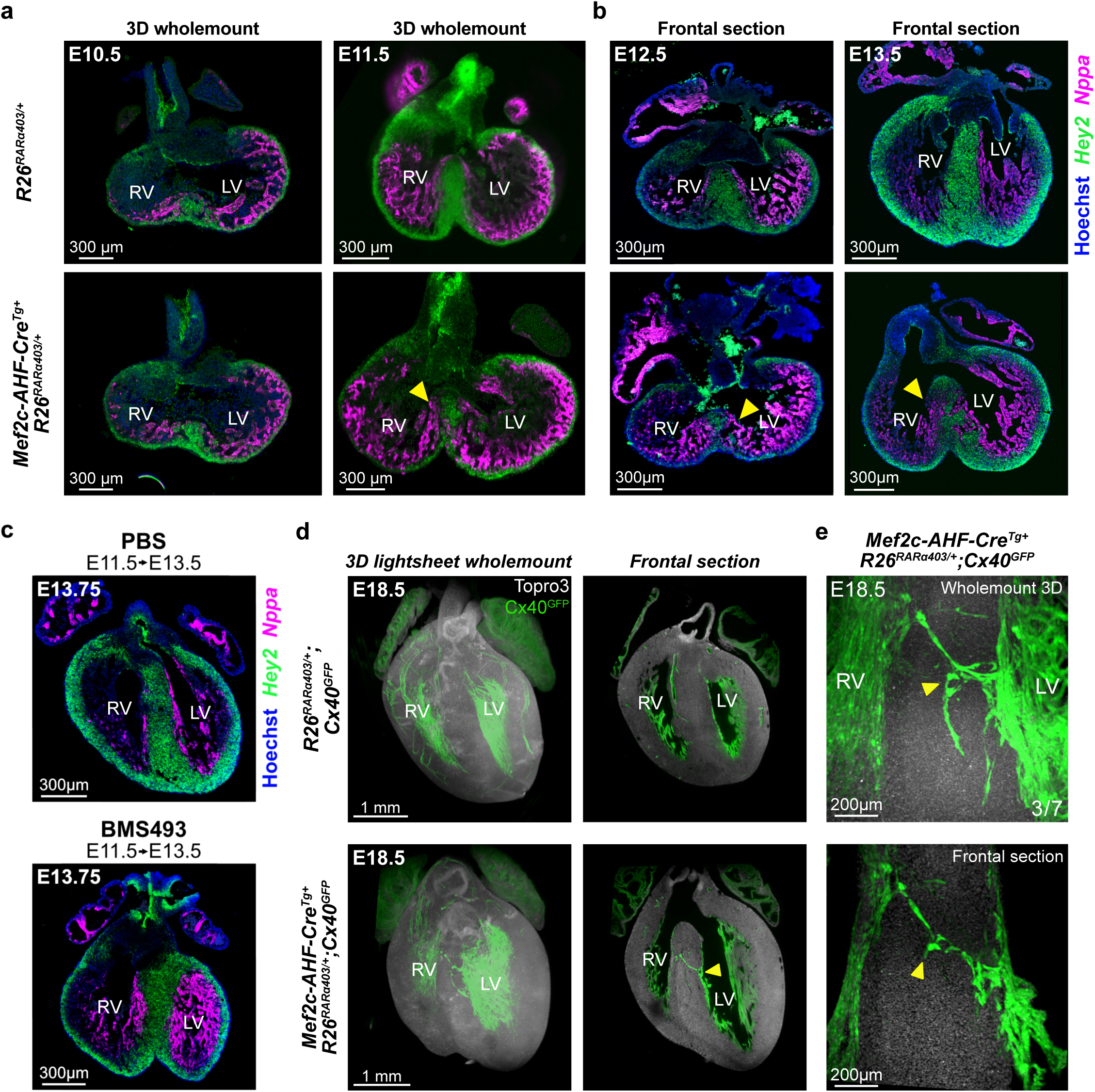
Expression of *RARα403* in the *Mef2c-AHF-Cre* genetic lineage leads to a trabeculated core at a stage preceding emergence of the bifid apex phenotype. **(a)** RNAscope labelling at E10.5 and E11.5 showing the expression of a trabecular gene, *Nppa* (magenta), and a compact myocardial gene, *Hey2* (green) in 2D planes of 3D light sheet images of *R26^RARα403^* and *Mef2c-AHF-Cre^Tg+^*; *R26^RARα403^* hearts. **(b)** RNAscope labelling of *Nppa* and *Hey2* transcripts at E12.5 and E13.5 in frontal paraffin sections of *R26^RARα403^* and *Mef2c-AHF-Cre^Tg+^*; *R26^RARα403^* hearts. Ectopic trabecular labelling is observed from E13.5 in the mutant septum (yellow arrowheads). **(c)** RNAscope labelling of *Nppa* (magenta), and *Hey2* (green) in 2D planes of E13.75 hearts collected after injection of the pan-RAR inverse agonist BMS493 on 3 consecutive days from E11.5 to E13.5. These hearts are bifid, but do not display ectopic trabeculation in the septum. **(d)** Wholemount (left) and frontal section (right) view of E18.5 cleared lightsheet imaged *R26^RARα403^*; *Cx40^GFP^* and *Mef2c-AHF-Cre^Tg+^*; *R26^RARα403^*; *Cx40^GFP^* hearts. Yellow arrowheads indicate ectopic conduction cells spanning the septum (observed in 3/7 mutants). **(e)** High resolution magnification of spanning conduction cells in the *Mef2c-AHF-Cre^Tg+^*; *R26^RARα403^*; *Cx40^GFP^* IVS shown in (d). IVS, interventricular septum.

*Gja5,* encoding Connexin 40, is expressed in trabecular myocardium and later in the ventricular conduction system ^49^. Analysis of an *Cx40(Gja5)^GFP^*allele revealed expanded expression in the septum of *Mef2c-AHF-Cre^Tg+^;R26^RARα403^* hearts at E12.5 (Fig. S8e). In E18.5 hearts analysed by lightsheet microscopy ectopic GFP labelling was observed in the central region of the septum in 3/7 *Mef2c-AHF-Cre^Tg+^;R26^RARα403^* hearts (Fig. 5d,e). These GFP positive cells are negative for CD31, a marker of endothelial cells, where *Gja5* is also expressed, suggesting that they correspond to ectopic conductive cardiomyocytes (Fig. S8f).

The trabeculated nature of the *Mef2c-AHF-Cre^Tg+^;R26^RARα403^*IVS prior to onset of the ventricular cleft phenotype was further confirmed by lightsheet imaging of hearts expressing a ubiquitous mGFP reporter gene between E10.5 and E12.5 (Fig. S9a). While by E12.5 all wild-type hearts have a compacted septum, some early wild-type E11.5 hearts were observed to have a partially non-compacted septal core (Fig. S9b; Video S5). Consistent with normal patterning of the IVS region in *Mef2c-AHF-Cre^Tg+^;R26^RARα403^* hearts, the septal primordium at E10.5, and at a region at the top of the septum at E11.5 and E12.5 (arrowheads in Fig. S9a), appeared unaffected. ScRNA-seq subclustering of myocardial cells revealed a population of *Hey2^+^* compact myocardial cells that did not change in relative cell number between control and mutant hearts; analysis of genes enriched in this subcluster revealed that this subpopulation expresses the Netrin 1 receptor encoding gene *Dcc* (Fig. 6a; Table S2). RNAscope staining revealed that *Dcc* labels a population of myocardial cells at the top of the septum that is present in *Mef2c-AHF-Cre^Tg+^;R26^RARα403^* hearts (Fig. 6b,c; Video S6). Moreover, *Dcc* is expressed in the septum primordium at E10.5 (Fig. S9c). The *Dcc* expressing RA-independent region may correspond to the inlet or non-folding region of the IVS^15, 19^. Together these results reveal sequential steps in muscular IVS morphogenesis: an early RA-independent step generating the septal primordium and subsequently the top of the compact septum, followed by RA-dependent compaction and fusion steps in the central and apical regions of the IVS.

**Figure 6.**
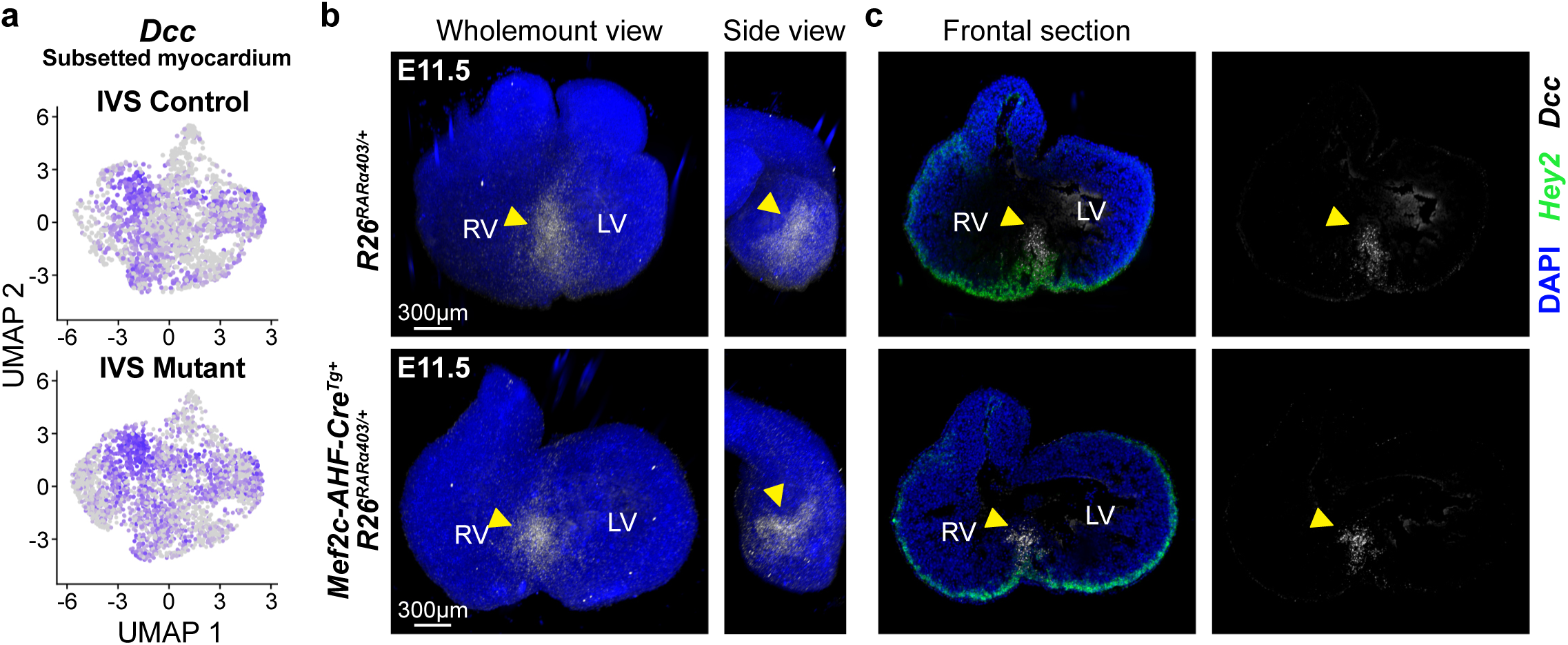
scRNA sequencing analysis identifies *Dcc* as a marker of a septal population not regulated by retinoic acid signalling. **(a)** Expression of *Dcc* in *R26^RARα403^* control and *Mef2c-AHF-Cre^Tg+^*; *R26^RARα403^* mutant subsetted myocardium. **(b)** Wholemount RNAscope labelling of *Dcc* (white) and *Hey2* (green) in E11.5 *R26^RARα403^* and *Mef2c-AHF-Cre^Tg+^*;*R26^RARα403^* hearts shown in wholemount views. **(c)** Frontal section views of the hearts in (b). *Dcc* expression labels a RA-independent domain at the top of the muscular septum, corresponding to the inlet septum. LV, left ventricle; RV, right ventricle.

### Retinoic acid signal reception modulates myocardial signalling

In order to further characterize RA-dependent gene expression in compact myocardium at E11.5, we exploited the fact that left ventricular cells from *Mef2c-AHF-Cre^Tg+^;R26^RARα403^* hearts do not express *Mef2c-AHF-Cre* and thus do not express *RARα403*. They can thereby serve as an internal control to identify DEGs. We subsetted the compact myocardial cluster of control and mutant cells and clustered them into left- and right ventricular cardiomyocytes, demarcated by *Tbx18* and *Pitx2* expression respectively ^4, 50^ (Figs. 7a and S10a,b). Validating this approach, we found that *Cre* transcripts were detected only in the mutant sample, primarily in the right ventricular cell cluster (Fig. 7b). As anticipated, we identified a higher number of DEGs in the right ventricular cluster when comparing mutant and control cells than in the left ventricular cluster; these included cardiac maturation and signalling related genes (Fig. S10c-f). Importantly, we observed right ventricular-specific downregulation of the known RA targets *Rarb* and *Mdk* ^39, 51^ (Figs. 7c and S10f), consistent with impaired RA signal reception.

**Figure 7.**
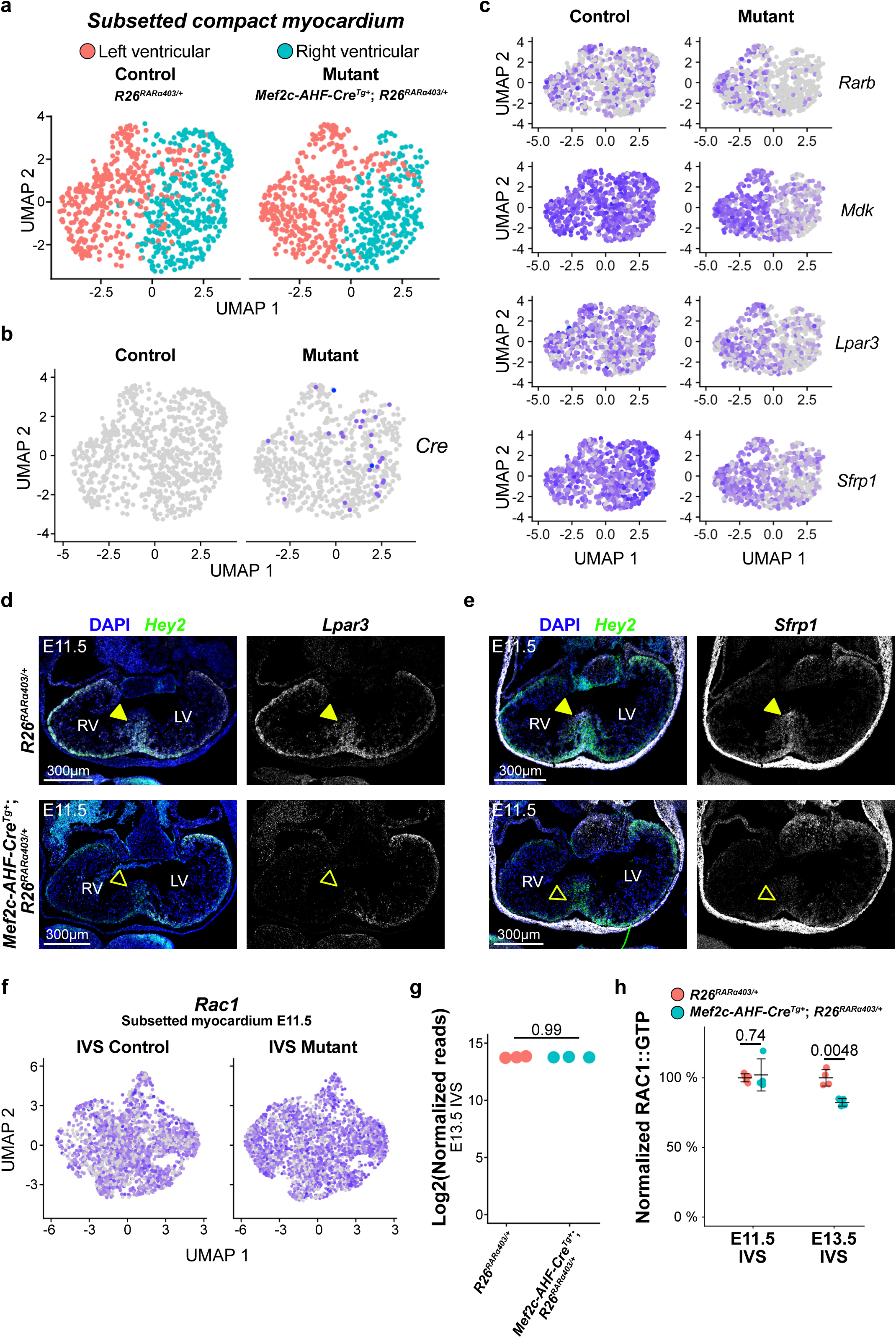
Single cell transcriptomic analysis reveals altered expression of signalling components and RAC1 activation on downregulation of retinoic acid signal reception. **(a)** UMAP of compact myocardial cells from *R26^RARα403^*and *Mef2c-AHF-Cre^Tg+^*;*R26^RARα403^* IVS subsetted and -clustered into left- and right ventricular compact myocardium. **(b)** Feature plot showing *Cre* expression in left- and right ventricular compact myocardial cells from control and mutant hearts. *Cre* is expressed only in mutants and predominantly in right ventricular cardiomyocytes. **(c)** Feature plots of selected DEGs. RA target genes *Rarb* and *Mdk* are downregulated only in RV compact myocardial cells in the mutant. Similarly, the genes encoding the lysophosphatidic acid receptor *Lpar3* and the modulator of Wnt signalling *Sfrp1* are downregulated in RV but not LV cells. **(d, e)** RNAscope ISH showing *Lpar3* (d) and *Sfrp1 (e)* expression in sections of E11.5 *R26^RARα403^* and *Mef2c-AHF-Cre^Tg+^*;*R26^RARα403^* embryos. Full arrowheads point to expression in the septum, while empty arrowheads in mutants indicate loss of expression. **(f)** Feature plot showing the expression of *Rac1* in control and mutant subsetted myocardium at E11.5. **(g)** Normalized read counts of *Rac1* transcript in bulk RNA sequencing data of E13.5 control and mutant IVS; P-value (adjusted p-values, DEseq2, n = [3 controls; 3 mutants]). See figure 8. **(h)** Quantifications of relative GTP-bound activated RAC1 levels by G-LISA (normalized to controls) of microdissected and pooled E11.5 and E13.5 IVS regions; P-value (Welch’s t-test) is shown. N [controls; mutants]: E11.5 [5;4], E13.5 [4;4]. IVS, interventricular septum; LV, left ventricle; RA, retinoic acid; RV, right ventricle; UMAP, Uniform Manifold Approximation and Projection.

Among the DEGs we identified a number of genes encoding molecules known to regulate intracellular signalling via the small GTPase RAC1. Intriguingly, second heart field specific deletion of *Rac1* or myocardial deletion of another Rho GTPase encoding gene *Cdc42* are the only other reported mouse mutants to our knowledge that also develop a bifid apex, among other severe cardiac defects^20, 52^. Among the candidate DEGs that have been shown to promote RAC1 activation are the lysophosphatidic acid receptor *Lpar3* and the soluble modulator of WNT signalling *Sfrp1*. RNAscope ISH at E11.5 revealed reduced *Lpar3* and *Sfrp1* transcript accumulation in the septal region and right ventricle of *Mef2c-AHF-Cre^Tg+^;R26^RARα403^* hearts (Fig. 7d,e). This result was further supported by wholemount RNAscope for *Sfrp1* followed by lightsheet imaging (Fig. S10g; Video S7), as well as by quantification of transcript levels, confirming significant downregulation of *Lpar3* and *Sfrp1* in the right ventricle and septum relative to the left ventricle (Fig. S10h,i). Upregulated DEGs included genes encoding inhibitors of RAC1 activity, such as *Srgap2* and *Arhgap* family genes (Fig. 7f; Table S1). We evaluated *Rac1* gene expression and protein activity in mutant hearts. While *Rac1* transcript levels are not significantly downregulated at E11.5 or E13.5 (Fig. 7f,g), the level of GTP-bound activated RAC1 is reduced by 19.2% in mutant compared to control hearts at E13.5 (p=0.0048), although unchanged at E11.5 (p=0.74) (Fig. 7h). These results point to mechanistic interactions between RA target genes and RAC1 activity in the etiology of the bifid ventricular phenotype.

### Additional roles of myocardial retinoic acid signal reception in ventricular maturation and coronary vasculature

To investigate the impact of decreased RA signal reception in fetal myocardium after the onset of the bifid ventricular phenotype we performed bulk RNA sequencing of microdissected IVS regions of *Mef2c-AHF-Cre^Tg+^;R26^RARα403^* and control *R26^RARα403^* hearts at E13.5 (Fig. S11a). 343 genes were found to be significantly upregulated and 317 significantly downregulated (Fig. 8a; Table S3). GSEA analysis revealed multiple changed gene sets, including downregulation of terms related to myocardial maturation and vascularization (Figs. 8b and S11b; Table S4). These results reveal broader roles of myocardial RA signal reception on ventricular development, in addition to the regulation of septal morphogenesis. Defects in myocardial maturation were validated in *Mef2c-AHF-Cre^Tg+^;R26^RARα403^* hearts. *Tnni2*, encoding fast Troponin I, a regulator of cardiac muscle contraction, is one of the most significantly downregulated genes identified by bulk RNA-seq at E13.5. RNAscope ISH at E12.5 revealed that *Tnni2* is expressed in a small region of the free left ventricle close to the ventricular base as well as in the compact myocardium of the right ventricular free wall and the right side of the IVS (Fig. 8c). In *Mef2c-AHF-Cre^Tg+^;R26^RARα403^* hearts, *Tnni2* expression is lost in the right ventricle and septal region but maintained in the small region of the left ventricle unlabelled by the *Mef2c-AHF-Cre* lineage (Fig. 8c). This is consistent with a role of RA signal reception in promoting myocardial maturation. In order to explore the kinetics of the differentially expressed genes, the ventricular septal regions of E11.5 to E14.5 hearts were collected (Fig. S11c) for qPCR of selected genes. *Tnni2* (Fig. 8d) was significantly downregulated at all stages, while upregulation of the endocardial gene *Nrg1* was only observed at E13.5 and E14.5 (Fig. 8e). We confirmed the kinetics of *Nrg1* expression by RNAscope hybridization at E11.5 and E13.5; increased endocardial labelling in the mutant septum is associated with greater trabecular contributions (Fig. S11d, e). In order to evaluate coronary vascularization, we characterized control and mutant E17.5 hearts carrying a *Cx40^GFP^* allele expressed in coronary artery endothelium (Fig. S11f). Segmentation of the coronary arteries revealed defects throughout the coronary tree (Fig 8f). Quantification revealed defects in extension of the right and septal coronary arteries in the apical region of the heart (Fig. 8g,h). Genetic lineage tracing confirmed that the coronary arteries are not derived from *Mef2c-AHF-Cre* expressing cells (Fig. S11g), suggesting that these effects are indirect. These results extend prior evidence for the importance of RA signalling in coronary vessel development ^38^.

**Figure 8.**
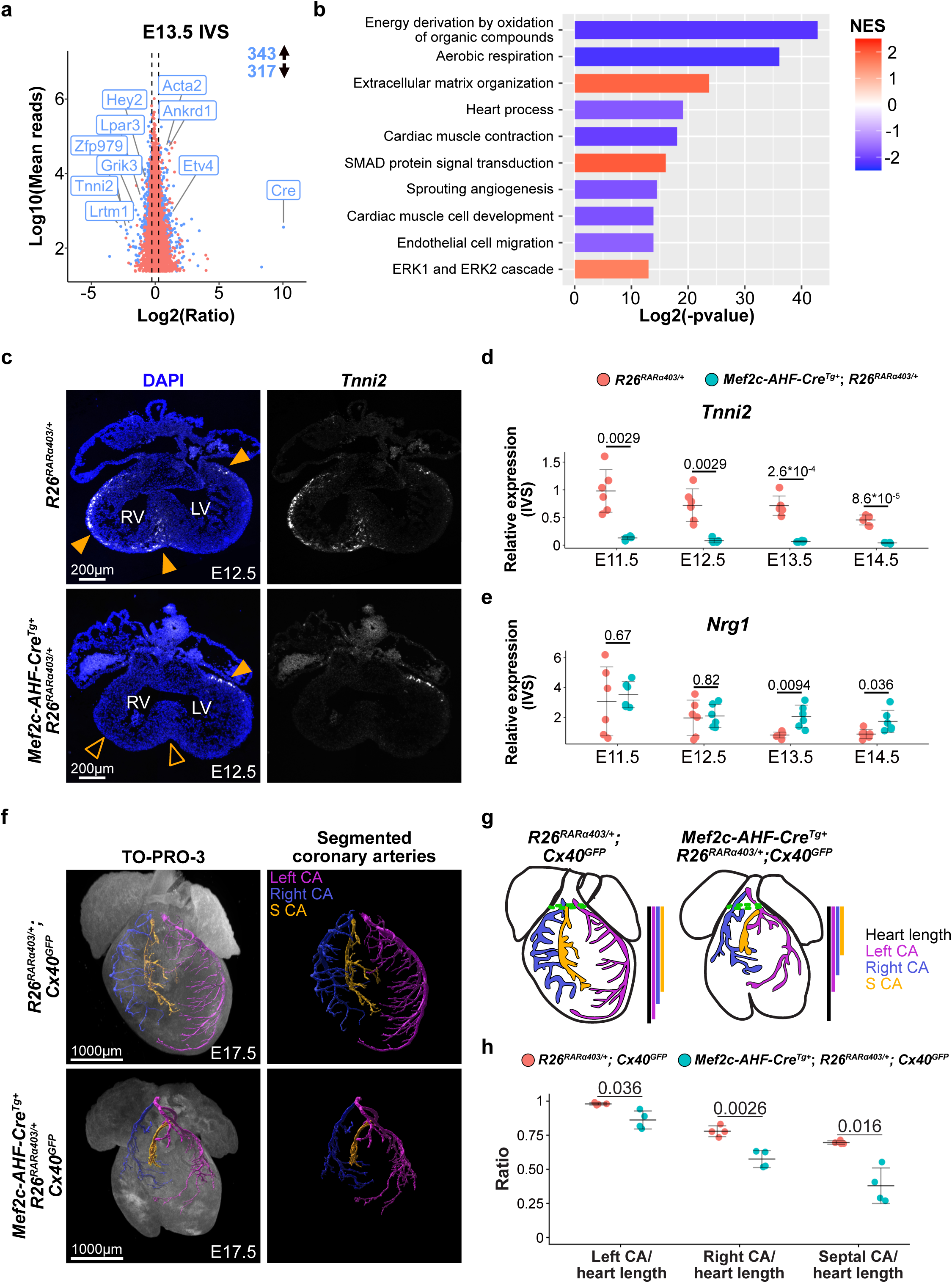
Downregulation of retinoic acid signal reception in myocardium leads to vascularization and differentiation defects in fetal hearts. **(a)** MA plot of relative gene expression levels in bulk RNA isolated from *R26^RARα403^* and *Mef2c-AHF-Cre^Tg+^*;*R26^RARα403^* IVS at E13.5. Non-significant differential expression is marked in red and significant differential gene expression in blue (Benjamini and Hochberg adjusted p-value < 0.05, DEseq2, n = 3 controls; 3 mutants). **(b)** Selected up and downregulated GSEA terms. Data is available in supplementary Table S4. **(c)** RNAscope for *Tnni2* in frontal sections of E12.5 *R26^RARα403^* and *Mef2c-AHF-Cre^Tg+^*;*R26^RARα403^* hearts. Full arrowheads indicate zones of expression, while empty arrowheads indicate zones that have lost expression in mutants. N = 3 control, 3 mutants. **(d, e)** Relative expression of *Tnni2* (d) and *Nrg1* (e) in micro-dissected IVS at stages E11.5 to E14.5 obtained by RT-qPCR. P-values (Welch’s t-test) are shown on figure. N for all stages = 6 controls, 6 mutants. **(f)** Segmented coronary arteries generated from wholemount lightsheet imaging of *R26^RARα403^*;*Cx40^GFP^*and *Mef2c-AHF-Cre^Tg+^*;*R26^RARα403^*;*Cx40^GFP^* hearts at E17.5, based on GFP expression shown in Fig S11C. **(g)** Measurement strategy to quantify length of apical extension of left-, right- and septal coronary arteries in hearts shown in (f). The dotted green line marks the border between ventricle and the great vessels. **(h)** Ratio of coronary artery apical extension versus heart length. P-values (Welch’s t-test) are shown on figure. N for all measurements [4 control; 4 mutants]. CA, coronary artery; IVS, interventricular septum; LV, left ventricle; RV, right ventricle; S-CA, septal coronary artery.

## Discussion

Our study provides new insights into the complexity of IVS development, identifying three distinct steps in septal morphogenesis (Fig. 9). These include early RA-independent generation of the septal primordium, followed by temporally distinct compaction and fusion steps in the central and apical regions of the IVS that depend on RA signal reception in the myocardium.

**Figure 9.**
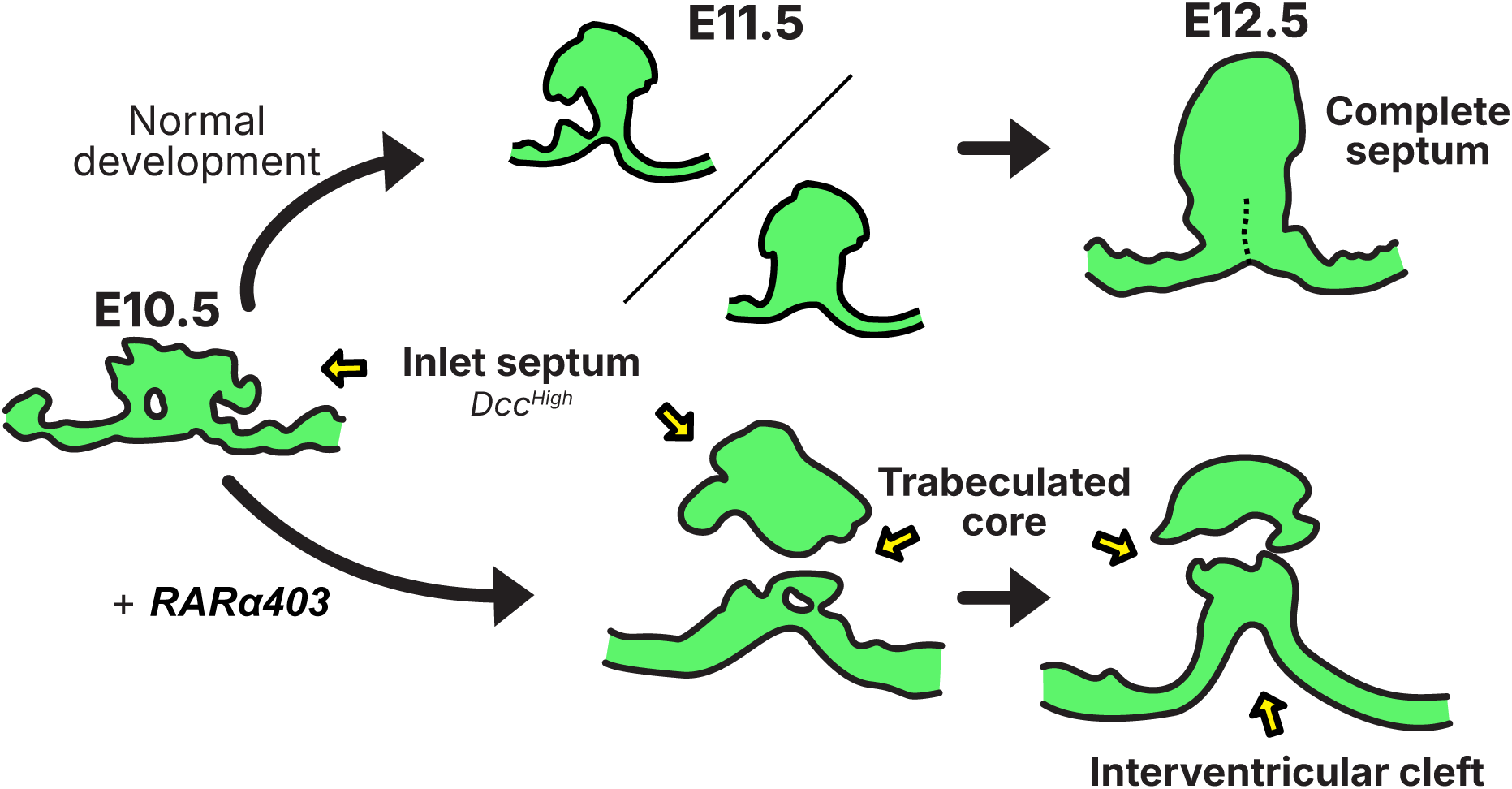
Cartoon illustrating retinoic acid dependent and independent steps of ventricular septal morphogenesis. Early patterning and formation of a *Dcc*-positive septal primordium is independent of myocardial RA signal reception, while formation of a compact central septal core at E11.5 and subsequent fusion of apposed septal proximal left and right ventricular walls in the apical region require myocardial RA signal reception. The dotted line represents the site of normal fusion of right and left ventricular walls.

The muscular IVS forms at the interface between cardiomyocytes derived from the first and second heart fields^4, 7^. An intersectional lineage has been identified by early expression of the *Mef2c-AHF* enhancer active in the SHF and the FHF regulator TBX5^7^. Subsequent downregulation of the SHF program in these interface cells defines a sharp boundary in the developing heart and the site of septum development^7, 42^. Indeed, elegant experiments have shown that TBX5 is required for septum formation in precisely this intersectional lineage that acts as a compartment boundary to regulate septation^8, 10^. Genetic tracing experiments suggest that these cells are derived from the earliest differentiating cardiomyocytes^7, 48^. Impaired RA signal reception in ventricular myocardium does not impact early patterning of the septal primordium. Indeed, the overlap between the *Mef2c-AHF* lineage and *Tbx5* expression and formation of the septal primordium occur normally in *Mef2c-AHF-Cre^Tg+^;R26^RARα403^* hearts. Our single cell profiling identified the gene encoding the Netrin receptor DCC as a marker of the early septal primordium; intriguingly, misexpression of *Netrin1* has recently been associated with septal defects in *Tbx5* mutant embryos^8^. Although labelled by *Mef2c-AHF-Cre*, this part of the septum does not depend on myocardial RA signal reception, supporting prior anatomical evidence for a distinct inlet septal component^15, 19^.

The epicardium is an established site of *Aldh1a1/2* expression and RA production, and RA has been shown to be important for normal ventricular development, in particular for growth of the compact myocardial layer, as initially demonstrated by the phenotype of mice lacking the RAR co-receptor RXRA^26, 29, 30^. Conditional genetic experiments suggested that the effect of RA on the myocardium was indirect, and mediated by secondary epicardial signalling events^31, 32, 34–36^. However, a recent study using a *RARE-CreERT2^Tg+^*transgene demonstrated that RA also signals directly to adjacent compact myocardium from mid-gestation, although the role of such direct signalling was not identified^41^. Our results confirm that compact myocardium responds directly to RA and show that downregulating RA signal reception specifically in SHF-derived ventricular myocardium leads to defects in septal morphogenesis, as well as in myocardial maturation and likely indirect effects on coronary vasculature. Single cell transcriptomics revealed that RA target genes such as *Rarb* and *Mdk* are downregulated in right ventricular cardiomyocytes in *Mef2c-AHF-Cre^Tg+^;R26^RARα403^* hearts, as is a *RARE-lacZ* reporter transgene. Together these results demonstrate that RA signal reception in myocardium is required for IVS morphogenesis. We observe ectopic trabecular myocardium in the forming septal region of *Mef2c-AHF-Cre^Tg+^;R26^RARα403^* hearts at E11.5, prior to quantifiable emergence of the interventricular cleft phenotype. This is similar to the expanded trabecular myocardium reported in *Rxra* null hearts^26^ and suggests a role for myocardial RA-signal reception in compaction of the septal core that may overlap with indirect RA inputs via the epicardium.

*Mef2c-AHF-Cre^Tg+^;R26^RARα403^* hearts develop a deep interventricular cleft and bifid ventricular phenotype. Our results support a folding model for normal septal development by which right and left ventricular walls converge in the septal region during ballooning morphogenesis^19^, followed by RA-dependent fusion to generate the apical muscular septum. While *Mef2c-AHF-Cre^Tg+^;R26^RARα403^* and *Rxra* null hearts both show an expansion of trabecular myocardium in the septal primordium at E11.5, fusion of the septal proximal left and right ventricular walls appears to be unaffected in *Rxra* mutant hearts. Moreover, this phenotype is not observed in other mutants with impaired development of the compact myocardial layer or hypertrabeculated hearts, identifying a previously undescribed RA-dependent step in ventricular development. Activating RARα403 using Cre lines expressed in different ventricular myocardial populations revealed that RA signal reception is required for apical fusion in cells that contribute to the core of the muscular septum. In addition, pharmacologically blocking RA signalling *in utero* during septal development results in an apical cleft without ectopic trabecular gene expression. Emergence of an interventricular cleft is associated with elevated expression of *Tbx18*, normally observed on the left side of the muscular septum, in the septal proximal left ventricular wall. Prior evidence for a sharp transcriptional and clonal boundary running along the length of the muscular IVS is consistent with cells on the left and right sides of the septum being initially separate and becoming apposed by infolding during septum morphogenesis^4, 6, 7^. Experiments in avian embryos suggest that epicardial cells becomes internalised in the septal core, coherent with juxtaposition of the epicardial surface of left and right septal proximal walls^19^ and epicardial ablation experiments indicate an essential role for the epicardium in septal morphogenesis^53^. Furthermore, trabecular gene expression analysis and genetic lineage studies have shown that the core of the septum shares the compact myocardial identity of the outer ventricular layer, consistent with an infolding model^12, 49^. Differences in the distribution of *RARE-CreERT2^Tg+^ Cre* expressing cells versus the RARE*-CreERT2^Tg+^* genetic lineage point to compact myocardial cell colonization of the septal core by E12.5. In summary, our temporal and spatial analysis of RA requirements for septal morphogenesis indicate that progressive infolding of the ventricular walls is followed by zipper-like RA-dependent fusion during formation of the muscular IVS.

A bifid ventricular apex is rare among mouse cardiac phenotypes. However, a similar phenotype has been observed in hearts in which *Rac1* is deleted in the SHF using *Mef2c-AHF-Cre*^20^. These mice have a range of severe cardiac defects including a bifid ventricular apex, implicating small Rho GTPase activity in septal morphogenesis. This phenotypic overlap suggests that RA-dependent signalling may also impact on small GTPase regulated processes such as cell migration during septal morphogenesis. Our single cell transcriptomic analysis identified a number of differentially expressed genes in septal myocardium of *Mef2c-AHF-Cre^Tg+^;R26^RARα403^* hearts encoding signalling molecules that potentially modulate RAC1 activity, including LPAR3 and SFRP1. Moreover, we observed reduced active RAC1 in microdissected septal tissue from mutant hearts at E13.5, but not E11.5, suggesting that myocardial RA signalling may act upstream of RAC1 activity during ventricular septation. Defects in myocardial maturation and development of coronary vasculature may also contribute to the apical cleft phenotype in *Mef2c-AHF-Cre^Tg+^;R26^RARα403^* hearts. Abnormal cardiomyocyte fibre development could, for example, lead to a failure to bridge right and left ventricular myocardium at the cardiac apex, and reduced vascular development impact signalling in the subepicardial region as right and left ventricular walls converge. Both excess and reduction of RA signalling have been shown to lead to reduced vascular coverage and impaired recruitment of epicardially derived smooth muscle cells ^38^. However, neither these hearts, nor fetal hearts lacking intraventricular coronary arteries, such as *Cxcl12* null mice^54^, have been reported to have a bifid ventricular apex. Future experiments will evaluate the potential contribution of these mechanisms, as well as RAC1 activation, to normal development of the muscular ventricular septum downstream of RA signalling to myocardium.

Finally, we consider the relevance of our findings for the origins of congenital heart defects and evolution of ventricular shape. While the apical cleft phenotype represents a defect in muscular septal morphogenesis there is no internal communication between the two ventricles, as in a membranous or muscular VSD^3^. In contrast, expanded trabecular myocardium in the central region of the septum may contribute to ectopic conductive cells and muscular VSDs. Similarly to the situation in the mouse embryo, the apex of the normal developing human heart is also transiently bifid, resolving on completion of septal morphogenesis. Rare human hearts with a ventricular cleft have been reported, either in isolation or associated with other defects, including conotruncal defects and sudden cardiac death, with unknown physiological consequences ^55, 56^. A deep interventricular cleft is observed in the hearts of diving mammals of the order Sirenia, such as the manatee and dugong ^57^; in such species uncoupling left and right ventricular contraction may optimize cardiac function on diving with apnea. Future efforts towards development of a postnatal mouse model will provide insights into this question. Furthermore, mutations in the RA signalling pathway may have contributed to changes in ventricular shape during mammalian evolution.

## Supporting information

Bonnelykke et al Supplementary Figures

## Acknowledgements

We are grateful to members of the Kelly lab, including Cemal Tastan, Harshit Pateria and Gaetano D’Amato and the following colleagues for discussion and input: Richard Harvey, Irfan Kathiriya, Fabienne Lescroart and Bjarke Jensen. We thank the IBDM and MMG imaging and animal platforms, Mai Huang (Turing Centre for Living Systems), the Genomics and Bioinformatics Platform (GBiM: Camille Humbert, Christel Castro and Valerie Delague), Inserm U1251, Marseille Medical Genetics Unit and Sebastien Dupichard (Zeiss).

This work was supported by the French Académie des Sciences, the Agence Nationale pour la Recherche (Heartbound ANR-22-CE13 and Heartist ANR-23-CE13-001 projects) the Fondation Leducq (Transatlantic Network of Excellence 15CVD01), the Fondation pour la Recherche Médicale (EQU202503020001), the Australian Research Council (Discovery Project grant DP210102134), the France-BioImaging/PICsL infrastructure (ANR-10-INSB-04-01) and the Turing Centre for Living Systems (France 2030, the French Government program managed by the French National Research Agency (ANR-16-CONV-0001) and the Excellence Initiative of Aix-Marseille University A*MIDEX).

## Author contributions

T.H.B., C. Coulon, R.S., M.C., C. Cortes, C.R., D.S., D.M., C.D., L.M. S.Z. and R.G.K. performed experiments; T.H.B., C. Coulon, R.S., M.C., C. Cortes, D.S., L.M., G.D., S.Z. and R.G.K analysed results; T.H.B. and R.G.K. wrote the manuscript; all authors reviewed and edited the manuscript; R.G.K. and T.H.B supervised the work and R.G.K. acquired funding.

## Material and methods

### Ethics statement

Animal experimentations were performed in strict accordance to the European Community Directive on the protection of animals used for scientific purposes (EU Directive 2010/63). Specific approval for all procedures presented were approved by the ethics committee of the IBDM SBEA and by the French Ministry of Research (APAFIS #44241-2023072110393026 v4) besides those involving the *RARE-CreERT2^Tg+^* line, which were performed under APAFIS #46272-2023121214179595 v3. Animal care, maintenance and husbandry in the CNRS-IBDM Animal Facility (G-13-055-21) fulfilled the animals’ welfare needs.

### Mice

The *Mef2c-AHF-Cre^Tg+^*, *R26^RARα403^*, *R26^mTmG^*, *RARE-lacZ^Tg+^*, *Z/EG^Tg+^*, *RARE-CreERT2^Tg+^*, *Mlc2v^Cre^*, *Acta2^CreERT2^*, *Lyz2^Cre^, R26^tdTom^*^ato^, *R26^RYFP^* and *Cx40^GFP^* mouse lines have previously been reported^9, 31, 39, 41–43, 45, 47, 58–61^. Mice were maintained on a mixed CD1 and C57Bl/6 background and genotyped by PCR using primers listed in Table S1. The *R26^mGFP^* constitutive expression mouse line was generated by crossing *R26^mTmG^* males with *Mef2c-AHF-Cre^Tg+^*females to achieve germline *Cre-loxP* recombination. Recombination was validated by PCR genotyping using the three primer 5’-CAACGTGCTGGTTATTGTGCT-3’, 5’-GCTCAGGTAGTGGTTGTCGG-3’ and 5’-ACCTTGTAGATCAGCGTGCC-3’ (537bp and 1263bp products for no recombination, 889bp product for recombination). Embryonic day (E) 0.5 was defined as noon of the day of the vaginal plug. Animals had food and water ad libitum and were housed in ventilated cages containing nesting material at room temperature of 22 ± 3°C, relative humidity between 45-65% and a 12h light-dark cycle.

### Tamoxifen and 4-OH tamoxifen injections

Tamoxifen (Sigma Aldrich, T5648) was dissolved in corn oil (Sigma-Aldrich, C8267-500ML) to a concentration of 20mg/ml. 200µl of this solution was injected intraperitoneally into pregnant females. For 4-OH tamoxifen injections, 4-OH tamoxifen (Sigma-Aldrich, H7904-25MG) was diluted to a 10mg/ml concentration in a 1:1 dilution of ethanol (VWR Chemicals, 20821.296P) and Cremophor EL (Millipore, 238470). This solution was diluted prior to injection to a 3mg/ml concentration in 1xPBS (Gibco, 14190-169) and 175-200µl was injected intraperitoneally (0.525-0.6mg total).

### BMS493 injections

BMS493 (Sigma-Aldrich, B6688) was diluted in Dimethyl-sulfoxide (DMSO) to a concentration of 10mg/ml and then further diluted in 1x PBS (Gibco, 14190-169) to a final concentration of 2mg/ml. This solution was injected intraperitoneally at a volume corresponding to 10mg/kg into pregnant females. DMSO:PBS (1:4) was used for injections in controls.

### Embryo collection and external heart imaging

Pregnant females were euthanized by cervical dislocation and embryos were dissected in 1x PBS (Eurobio Scientific, CS0PBS01-08) treated with diethyl pyrocarbonate (DEPC, Sigma-Aldrich, D5758-100ML). Depending on the experiment, either full embryos or hearts were collected. Hearts used for size- or length quantifications were treated with 250mM ice cold KCl for 5-10 minutes to arrest them in diastole. Fixation of embryos and hearts were done for 24 hours at 4°C in 1X PBS containing 4% paraformaldehyde (Election Microscopy Sciences, 15714) followed by washes in 0.1% PBT (1X PBS containing 0.1% Tween-20 (Euromedex, 2001-A)). External heart images for heart length and bifid cleft measurements were acquired on a Leica S9i stereo microscope or a Zeiss AxioZoom V16 microscope equipped with an Axiocam 512 colour camera. For long-term storage, paraffin embedding or wholemount RNAscope *in situ* hybridization, embryos and hearts were dehydrated into 100% methanol (VWR Chemicals, 20847.295).

### Measurements of bifid cleft ratio of external heart images

Measurements were performed using Fiji (c2.16.0). To measure the relative heart- and bifid cleft length, a line was drawn between the most apical point of each ventricle. From this line, two perpendicular lines were drawn: one going to the base of the ventricular outlet (heart length) and one going to the centre of the ventricular cleft (bifid cleft length).

### Histology - Formalin fixed, paraffin embedding (FFPE)

Dehydrated hearts and embryos were washed for 2 hours in xylene (VWR chemicals, 28975.291) and embedded in paraffin (Paraplast X-tra, Sigma-Aldrich, P3808-1KG). 10µm transverse sections of embryos or frontal sections of isolated hearts were mounted on Superfrost Plus Adhesion Microscope Slides (Epredia, J1800AMNZ) and stored at 4°C prior to staining.

### Antibodies

The following antibodies were used: Mouse α MF20 (targeting MYH1E) (1:50, DHSB), rat α CD31 (1:200, Dianova, DIA-310), rabbit α Ki67 (1:500, Abcam, ab15580), rabbit α Cleaved Caspase-3 (Asp175) (1:500, Cell Signaling Technology, #9661), mouse α TBX5 (1:100, Santa Cruz, sc-515536), chick α GFP (1:500, Avés Labs, GFP-1020), Alexa Flour 488 donkey α chicken (1:500, Jackson ImmunoResearch, #703-545-155), Alexa Flour 488 donkey α mouse (1:500, ThermoFisher Scientific, A21202), Alexa Flour 568 donkey α mouse (1:500, Life Technologies, A10037), Alexa Flour 647 donkey α rabbit (1:500, ThermoFisher Scientific, A31573), Biotin donkey α rat (1:200, Jackson ImmunoResearch, #712-065-153) and Biotin donkey α rabbit (1:200, Jackson ImmunoResearch, #711-065-152). For signal amplification, HRP-streptavidin (1:300, Revvity, NEL750001EA) and TSA-Cy3 (1:300, Akoya Biosciences, TS-000202) or TSA-Vivid Fluorophore 650 (1:1000, ACD Bio-Techne, #323273) in 1x Amplification Diluent (Perkin Elmer, FP1135) were used.

### Immunofluorescent labelling of FFPE sections

Sections were permeabilized in 1x PBS containing 0.2% Triton X-100 for 20 min, followed by blocking in TNB blocking buffer (Roche Diagnostic, #11096176001) for 1 hour. Primary antibodies were incubated in TNB blocking buffer and incubated overnight at 4°C. Secondary antibodies coupled to fluorescent molecules were incubated in TNB blocking buffer for 1 h along with DAPI for nuclear labelling (1:1000, Invitrogen, 62248). Slides were mounted in Fluoromount-G (Southern Biotech, 0100-01) and imaged on a Zeiss AxioZoom V16 microscope equipped with an Axiocam 512 colour camera or a Zeiss LSM 880 confocal microscope.

### Wholemount X-gal staining and sectioning

Hearts were collected from *RARE-LacZ^Tg+^* embryos in 1x PBS and then fixed for 15 minutes at 4°C in 4% PFA. After three washes in PBS, hearts were incubated in X-gal staining solution overnight at 37°C [0.1% X-gal (Millipore, 4063-102), 2 mM MgCl2, 0.01% deoxycholate, 0.01% Nonidet P40, 5 mM potassium hexacyanoferrate (II) trihydrate (Sigma-Aldrich, P3289) and 5 mM potassium hexacyanoferrate (III) (Sigma-Aldrich, 244023)]. Staining was stopped by three washes in 1x PBS followed by fixation for 1 hour at 4°C (4% PFA). Selected hearts were then embedded in paraffin and sectioned in 8 µm sections. Both wholemount- and section images were acquired on a Zeiss AxioZoom V16 microscope.

### Immunofluorescence on cryosections

After fixation, hearts were washed in 1x PBS containing 0.1% Tween-20 (PBT 0.1%) and then washed in progressive (15%, 30%) sucrose baths in PBS before being incubated overnight in OCT (VWR Chemicals, 361603E). Samples were then transferred to new OCT on dry ice and oriented prior to sectioning. 10µm transverse sections were made using a Leica CM3050 S Cryostat, mounted on Superfrost Plus Adhesion Microscope Slides (Epredia, J1800AMNZ) and stored at -70°C. For immunofluorescent labelling, slides were washed three times in 1x PBS containing 0.05% Tween-20, followed by permeabilization for 15 minutes in 1x PBS containing 0.2% Triton X-100 (Euromedex, 2000-B) and signal bleaching in 1x PBS containing 3% H_2_O_2_ for 10 minutes. After three washes in PBT 0.05%, hearts were incubated for 1h in a blocking solution (1x PBS containing 2% bovine serum albumin (Sigma-Aldrich, A7030) and 0.05% Saponin (Millipore, 558255)) at room temperature. Slides were then incubated with primary antibodies diluted in blocking solution overnight at 4°C. After three washes in PBT 0.05%, sections were incubated with the secondary antibodies and Hoechst 33258 (1:1000) diluted in the blocking solution for 1 h at room temperature. Slides were mounted in Fluoromount-G (Southern Biotech) and imaged on a Zeiss LSM780 confocal microscope.

### Wholemount immunofluorescent staining

Hearts were washed in 1x PBS containing 0.1% Tween-20 (PBT) after fixation and put in TNB blocking solution (Roche Diagnostics, #11096176001) overnight at 4°C. Following this, hearts were incubated for 48 hours with primary antibodies in TNB blocking buffer at 4°C, washed thrice in PBT and incubated 24 hours with secondary antibodies and Hoechst 33258 (1/1000, Merck Sigma-Aldrich, 861405) (1:1000) at 4°C. After three PBT washes, the hearts were imaged on a Zeiss LSM 880 confocal microscope.

### Quantification of proliferation index in sections

3-5 immunofluorescent labelled sections per heart were imaged on a Zeiss LSM 880 confocal microscope using a 20x objective. Morphological quantification of the 2D immunofluorescence images were done using the Fiji software. IVS was selected as the ROI and different channels were separated using the tissue specific staining of the myocardium, endocardium, Ki67 and nuclei. Based on the threshold of the tissue specific staining, subsequent myocardial and endocardial masks were generated. Nuclei individualization for the ROI was done using the Stardist plugin in Fiji software followed by identification of Ki67^+^ and Ki67^-^ cells by opening Ki67 channel in tandem with the nuclear channel. Tissue areas, cell numbers, cell density and the proliferative index were quantified using the mask generated for the different tissues.

### RNAscope *in situ* hybridization

*In situ* RNAscope labelling’s were performed on both on FFPE sections and on whole mount hearts as described ^44, 62^, using the Multiplex Fluorescent v2 Assay (Biotechne, 313110), with the only change being that for whole mount samples, hearts were treated with 3% H_2_O_2_ (ACD Biotechne, 322381) for 30 minutes prior to RNAscope labelling. *Hey2* (404651-C1/C2), *Cre-O4* (546951-C2), *Cre-04* (546951-C3), *tdTomato* (317041-C3), *Tbx5* (519581-C2), *Tbx18* (515221-C1), *Tnni2* (1065311-C1), *Emcn* (548311-C1), *Nrg1* (418181-C3), *Dcc* (427491-C3), *Lpar3* (432591-C2) and *Sfrp1* (404981-C1) probes were used. Fluorescent signal amplifications were obtained using TSA Plus Cy5 (Akoya Biosciences, TS-000203), TSA Plus Cy3 (Akoya Biosciences, TS-000202) and TSA Plus Fluorescein (Perkin Elmer, FP1168). DAPI (Invitrogen, 62248) was used as nuclear counterstain in 1:500 concentrations in wholemount and 1:10,000 concentrations in FFPE sections. For FFPE sections, images were acquired on a Zeiss AxioImager Z1 Apotome1.2. For wholemount samples, samples were cleared in CUBIC R+(M)^63^ overnight at 4°C and then embedded into refractory index matched 2% agarose-CUBIC R+(M) (Promega, V2111) and imaged on a Zeiss Lightsheet 7 using a 5x / 0.16 focal clearing immersion detection objective.

### Quantification of ventricle wall thickness

To score the thickness of ventricular walls from E10.5 to E13.5 of *Hey2* and *Nppa* RNAscope in labelled sections, the InteredgeDistance v1.0.1 macro of Santosh Patnaik (available at: https://forum.image.sc/t/imagej-macro-to-measure-distance-between-two-lines-edges/42019) was used in Fiji (c2.16.0) with the adjustment to compute on average 50 lines to measure the average thickness. Using the *Hey2* signal, the outer- and inner side of the ventricles were traced and the average distance between these lines was calculated.

### Quantification of RNAscope signal ratio in sections

In Fiji, the compact myocardium of the left- and right ventricle were segmented using a *Hey2* labelling. The interventricular septum was also segmented. To generate a mask of signal positive and negative regions, an Otsu automatic thresholding was used on the left ventricular segmentation. This threshold was then applied to the whole image. For each segmented region, the ratio of the signal positive region versus the area was computed. For each sample, 2-4 sections were analysed.

### Activated RAC1::GTP quantification

The IVS of E13.5 samples were micro-dissected (Fig. S11A) in ice cold 1x PBS and flash frozen. To score the levels of RAC1::GTP, we used the Rac1 G-LISA GTPase Activation Assay kit (Cytoskeleton Inc., #BK126). In short, the IVSs were pooled (10-11 IVS at E11.5, 2 IVS at E13.5) and proteins extracted according to manufacturer’s recommendations. Protein levels were quantified using a Cell Density Meter (Fischer Scientific) and samples diluted to a 0.5 mg/ml concentration. All samples were performed in three technical replicates and quantified on a CLARIOstar Plus microplate reader (BMG Labtech). Technical replicates varying more than 10% from other replicates were excluded.

### Cubic clearing and lightsheet imaging

In this study, two clearing protocols were used: for E14.5 samples used for 2D image plane quantifications (Fig. 1G-H), sample clearing was obtaining using the CUBIC R1-R2 protocols^64^, while for all other experiments, sample clearing was obtained by the CUBIC L-R+(M) protocol^63^. In brief, samples were incubated 1-3 days in the lipid removal solutions (CUBIC R1 / CUBIC L) at 4°C followed by three washes and incubation with the nuclear marker TO-PRO-3 (1:500, Life Technologies, T3605) for 2-4 days in 0.1% PBT. Samples were then cleared in CUBIC R2 / CUBIC R+(M) at 4°C or room temperature for 2-4 days and then embedded into refractory index matched 2% agarose-CUBIC R2 / R+(M). Images were acquired on a Zeiss Lightsheet 7 using a 5x / 0.16 focal clearing immersion detection objective or a Clr Plan-Neofluor 20x / 1.0 focal clearing immersion detection objective (Refractory index: 1.53). 3D Images were visualized using Imaris and Imaris Viewer (v11.0.0, Oxford Instruments).

### Vibratome sectioning and antibody-labelling after clearing

Hearts were de-embedded from cubic-agarose by adding them to 1x PBS and heating them with mild shaking on a thermoshaker for 30 minutes at 65°C. Following three 1x PBS washes, the hearts were embedded in molds containing water with 8% sucrose and 3.5% agarose and placed on ice. Hearts were sectioned in 50 μm sections on a vibratome and placed in a 12-well plate containing a blocking solution. After 2 hours at room temperature with mild agitation, primary antibodies were added for overnight staining at 4°C. After 3 1x PBS washes, secondary antibodies were added for 3h at room temperature, followed by a three 1x PBS washes. Sections were mounted on a Superfrost Plus Adhesion Microscope Slides (Epredia, J1800AMNZ) and imaged on a Zeiss AxioImager Z1 Apotome1.2.

### Measurement on 2D planes of E14.5 cleared hearts

Images were oriented in 3D to find matching 2D planes on Imaris (v11.0.0). Lines were drawn manually on these optical sections to measure heart- and cleft length (as described above) and to measure apex to apex length. The outer curvature of the left ventricle was manually traced from the start of the AVC to the LV apex or the centre of the cleft to measure these distances.

### Measurement of the extent of GFP signal in *Z/EG^Tg+^* hearts

Images were oriented in 3D to find matching 2D planes using Imaris (v11.0.0). The ventricle width was measured by drawing a line from the widest points of the right- and left ventricles. The extent of the GFP signal was determined using the same point in the right ventricle and drawing a line to the furthest GFP^+^ point on the heart-width line.

### Segmentation of coronary arteries

Segmentation of coronary arteries was performed using Imaris software on 3D lightsheet images of 4 control (*R26^RARα403/+^*; *Cx40^GFP/+^*) and 4 mutant (*Mef2c-AHF-Cre^Tg+^*;*R26^RARα403/+^*; *Cx40^GFP/+^*) hearts from two separate litters that had been processed for CUBIC cleared imaging simultaneously. First, automatic segmentation was used to generate the overall shape of the coronary arteries, followed by manual correction, using the GFP signal. Afterwards, the left-, right- and septal coronary arteries were separated and a mask generated to visualize the network. Small vessels that did not connect to the main branches were excluded.

### Measurement of the apical extension of coronary arteries

Using matched 2D planes, heart length was measured for each heart by drawing a line between the cardiac apex and ventricular oultet. On this line, points were made that represent the projection of the furthest apical extension of each coronary artery. For each coronary artery apical extension was calculated as a ratio relative to heart length.

### RNA isolation

The ventricles of CD1 wild-type E13.5 hearts and the IVS of control (*R26^RARα403/+^*) and mutant (*Mef2c-AHF-Cre^Tg+^*;*R26^RARα403/+^*) E11.5-E14.5 hearts were micro-dissected in ice-cold 1x PBS (Gibco, 14190-169) and flash frozen in liquid nitrogen (for IVS dissections, see Figs. S7a and S11a,c). To isolate RNA, TRIzol-chloroform extraction (Invitrogen, 15596026) was performed followed by purification using the RNeasy Micro kit (QIAGEN, 74004). Extracted RNA was stored at -70°C.

### Reverse transcription quantitative PCR (RT-qPCR)

RNA concentrations were measured on a Nanodrop One (Thermo Scientific) and cDNA was generated using the iScript kit (Bio-Rad, #1708841) on RNA from single IVS (no samples were pooled). The amount of RNA used for reverse transcription ranged from 200ng for E11.5 IVS to 500ng for E14.5 IVS and E13.5 wild-type ventricles. RT-qPCR reactions were performed using PowerSYBR Green (Applied Biosystem, 2402654) on a CFX-96 system (Bio-Rad). *Hprt1* was used as a reference gene and cDNA from E13.5 wild-type ventricles was used as reference sample between different reactions. Primers are listed in Table S5.

### Bioinformatic analyses of published scRNA-seq data

Data from cardiac wild-type CD1 cells at E9.5-P9^46^ were downloaded from https://www.ncbi.nlm.nih.gov/geo/query/acc.cgi?acc=GSE193346 and restricted from E10.5 to E14.5 and the shown populations. It was visualized using Seurat v5.3.0^65^. For the FeaturePlot in Figure S5c, ventricular cardiomyocytes were subsetted and a new UMAP was generated. To highlight positive cells, the cells were plotted by order of expression by setting order = TRUE.

### Bulk RNA sequencing

For bulk RNA sequencing of the RNA extracted from E13.5 micro-dissected IVS samples, RNA quantity and -quality were measured using Agilent RNA 6000 Nano chips on a BioAnalyzer (Agilent). All RINs were ≥ 9.40. 250ng of RNA per sample was used for preparing mRNA libraries with the Kapa mRNA HyperPrep (Roche, #KR1352-v5.17) kit. Paired-end sequencing of 75 base pair reads was performed on an Illumina NextSeq 500 platform using equimolar amounts of libraries per sample. The mean sequencing depth per sample was 57.5 million reads.

### Bulk RNA analysis

FASTQ files were mapped to the mouse reference genome GRCm38/mm10 with the addition of sequences for *Cre* (NC_005856.1), *R26^RARα403^* (Nucleotide 1 to 1739 of NM_001024809.4) and *Sv40-polyA* using STAR v2.7.2b^66^, and bam files were indexed using Sambamba v0.6.6^67^. After mapping, the number of reads per feature was determined using Stringtie v1.3.1c^68^. Normalization and differentially expressed genes were calculated using DESeq2^69^ and p values were adjusted for multiple testing using the Benjamini-Hochberg procedure. Gene set enrichment analysis (GSEA) was performed by using the gseGO function from the R package clusterProfiler (v3.0.4) on the stat parameter from DEseq2 analysis^70^. The data set has been submitted to the ArrayExpress database (http://www.ebi.ac.uk/arrayexpress) under accession number: E-MTAB-16670.

### scRNA single cell preparation and sequencing

E11.5 IVS were microdissected and stored in ice cold 1x PBS (Gibco, 14190-169) containing 1% Fetal Bovine Serum (FBS) until completion of genotyping, upon which 10 control IVS and 8 mutant IVS were pooled by genotype. Cell dissociation was performed in a mix of collagenase IV (1:10, Worthington Biochemicals, #LS004186), Dispase (1:50, Worthington Biochemicals, #LS02100) and DNase I (500U/mL, Worthington Biochemicals, #LS002007) in 1x PBS by slow, continuous pipetting at 37°C. Dissociation was inhibited in 1x PBS containing 10% FBS. Cells were counted in a Countess 3 Automated Cell Counter (Invitrogen). The scRNA library was generated using the Chromium Next GEM Single Cell 3’ GEM kit (v3.1) on a Chromium Controller (10x Genomics) and sequencing was performed on a NovaSeq6000 (Illumina).

### scRNA analysis

FASTQ alignment and UMI counts quantifications were generated by Cell Ranger 8.0.1 (10x Genomics) using the mouse reference genome GRCm38/mm10 with the addition of sequences for *Cre*, *RARα403* and *Sv40-polyA* (see Bulk RNA sequencing analysis). Downstream analysis was performed in R (v.4.5.1) Corrections for ambient RNA was performed using soupX^71^ and doublets were predicted using scDblFinder^72^. Quality control, normalization, clustering, integration and visualization were done in Seurat (v5.3.0) using standard procedures^65^. Markers used for annotating clusters are shown in figure S7B-C. Red- and white blood cells were removed and all four major clusters were subsetted and reclustered. Cells from clusters with a high percentage of predicted doublets were removed. After quality control we obtained 5973 control cells (average 3830.0 genes and 13712.1 UMIs detected per cell) and 9923 mutant cells (average 3059.7 genes and 9227.4 UMIs detected per cell). Differences in cell type distribution between control- and mutant subsetted myocardium were tested by a simulated Fisher’s test (2000 iterations). To calculate differentially expressed genes, the Seurat function FindMarkers was used with the MAST statistical test^73^ (Table S1). To calculate the positive markers of the *Dcc^High^* compact myocardial cell cluster, the Seurat function FindConservedMarkers was used (Table S2). Mitochondrial-, ribosomal- and predicted genes were excluded. To subcluster left-and right ventricular compact myocardial cells, the cells were first subclustered and separated into control and mutant. Control cells were clustered alone and mutant cell identity was predicted based on control cell identity using the Seurat function FindTransferAnchors. Mutant cells with prediction scores < 0.65 were excluded. The data set has been submitted to the ArrayExpress database (http://www.ebi.ac.uk/arrayexpress) under accession number: E-MTAB-17365.

### Statistical tests and data visualisation

All measurements were taken from distinct samples and no data besides single outlier points in the RAC1 activation assay were excluded from analysis. Normal distribution was assumed for all experiments. All statistical tests were performed in R (v.4.5.1). Tests are indicated in the figure legends. Data was visualized using the ggplot2 package and available in the source data supplementary file.

## Supplementary figure legends

**Figure S1. Related to Figure 1. Quantification of altered ventricular septal morphology on expression of *RARα403* in the *Mef2c-AHF-Cre* genetic lineage. (a)** Wholemount view of cleared wholemount fluorescent lightsheet imaged *Mef2c-AHF-Cre^Tg+^*;*R26^mTmG^* hearts at E11.5, E12.5 and E14.5; frontal sections are shown in Fig. 1A. **(b)** Ventral brightfield images of *R26^RARα403^* and *Mef2c-AHF-Cre^Tg+^*;*R26^RARα403^* hearts at E10.5, E15.5 and E18.5. Black arrowheads indicate mutant interventricular clefts. **(c)** Ventral brightfield images of *Mef2c-AHF-Cre^Tg+^*;*R26^RARα403^* hearts at E18.5. 8/20 hearts display RV hypoplasia (arrowhead), while 2/20 hearts are enlarged. **(d)** Wholemount and frontal section views of TO-PRO-3 nuclei labelled cleared *R26^RARα403^*and *Mef2c-AHF-Cre^Tg+^*;*R26^RARα403^* hearts at E18.5. White arrowheads indicate the interventricular cleft in mutant hearts **(e)** Frontal and right sided views of wholemount fluorescent image of TO-PRO-3 nuclei labelled cleared *R26^RARα403^* and *Mef2c-AHF-Cre^Tg+^*;*R26^RARα403^* hearts at E14.5. Yellow dotted lines indicate the planes shown in Fig 1g; the white arrowhead indicates the interventricular cleft. **(f)** Additional measurements of E14.5 hearts based on 2D planes shown in Fig. 1g. Heart lengths are equivalent between controls and mutants, while the cleft size and distance between the RV and LV apex is larger in mutants. Means, standard deviations and P-values (Welch’s t-test) are shown. N [Control; Mutant]: Heart length and cleft length [10;11], Apex to apex distance [9;10]; IVS, interventricular septum; LV, left ventricle; RV, right ventricle.

**Figure S2. Related to Figure 1. No significant differences are observed in proliferation or apoptosis in *Mef2c-AHF-Cre^Tg+^*;*R26^RARα403^* hearts at E12.5**. **(a)** Confocal images of antibody labelled frontal sections of E12.5 *R26^RARα403^* and *Mef2c-AHF-Cre^Tg+^*;*R26^RARα403^* hearts labelled for myocardium (MF20, green), endocardium (CD31, red), active phases of cell cycle (Ki67, orange) and nuclei (DAPI, blue). **(b)** Quantifications of the septal regions of the images in (a) using filters to score myocardial and endocardial cells. Mutant septa have a smaller area, lower number of cells and display a tendency of reduced cell density. However, the ratio of Ki67+ cells (the proliferative index) is similar between control and mutants. Means, standard deviations and P-values (Welch’s t-test) are shown. N [3 Control; 3 Mutant]. **(c)** Fluorescent imaging of cleaved Caspase 3 (white) antibody labelled frontal sections of E12.5 *R26^RARα403^* and *Mef2c-AHF-Cre^Tg+^*;*R26^RARα403^* E12.5 hearts. Nuclei are labelled by DAPI (blue). Very few cleaved Caspase3^+^ cells are observed (yellow arrowhead) in both control and mutant ventricles. IVS, interventricular septum; LV, left ventricle; RV, right ventricle.

**Figure S3. Related to Figure 1. Reduced retinoic acid signalling in the right ventricle upon expression of *RARα403* in the *Mef2c-AHF-Cre* genetic lineage. (a)** Wholemount β-galactosidase staining of retinoic acid signalling reporter *RARE-LacZ^Tg+^* control and mutant hearts at E12.5. Red filled arrowheads point to transgene expressing regions that are absent in mutant hearts (empty arrowheads). **(b)** Histological analysis reveals that both the epicardium (black arrowheads) and the compact myocardium (magenta arrowheads) display β-gal^+^ cells in the right ventricle of control hearts. In *Mef2c-AHF-Cre^Tg+^*;*R26^RARα403^* hearts, labelling is observed in the epicardium but not in right ventricular myocardium (empty magenta arrowhead). N [control; mutants]: wholemount [7; 2], sections [4;2]. IVS, interventricular septum; LV, left ventricle; RV, right ventricle.

**Figure S4. Related to Figure 2. Retinoic acid signalling does not affect early patterning of the interventricular region. (a)** MIP of wholemount fluorescent image of lineage traced *Mef2c-AHF-Cre^Tg+^;Z/EG^Tg^*^+^ and *Mef2c-AHF-Cre^Tg+^;Z/EG^Tg+^;R26^RARα403^* hearts at E10.5 in ventral and dorsal views labelled for GFP (green) and TBX5 (red). White arrowheads point to GFP^+^; TBX5^+^ double positive regions. N [3 controls; 2 mutants]. **(b)** Frontal sections of wholemount fluorescent imaged, TO-PRO-3 nuclei labelled and cleared *Mef2c-AHF-Cre^Tg+^;Z/EG^Tg^*^+^ and *Mef2c-AHF-Cre^Tg+^;Z/EG^Tg+^;R26^RARα403^* hearts at stage E11.5. **(c)** Quantification strategy for measuring the extent of GFP signal in hearts from (B). **(d)** Quantifications of hearts from (b) using the strategy in (c). Means, standard deviations and P-values (Welch’s t-test) are shown. N [8 Control; 3 Mutant]. **(e)** RNAscope showing *Tbx5* transcript distribution in frontal sections of E13.5 *R26^RARα403^* and *Mef2c-AHF-Cre^Tg+^*;*R26^RARα403^* hearts. (**f)** RNAscope for *Tbx18* in frontal sections of E13.5 *R26^RARα403^* and *Mef2c-AHF-Cre^Tg+^*;*R26^RARα403^* hearts; the limit of *Tbx18* expression in the septal proximal left ventricular wall is indicated by arrowheads (n = 6). Abbreviations: LV, left ventricle; RV, right ventricle.

**Figure S5. Related to Figure 2. Retinoic acid signalling during ventricular septum formation. (a)** Frontal views of RNAscope labelled *Hey2* (green) and *Cre* (magenta) *RARE-CreERT2^Tg+^* hearts at E10.5 to E13.5, shown in high magnification in Fig. 2c. **(b)** Violin plots showing expression of the three aldehyde dehydrogenase encoding genes (*Aldh1a1/2/3*) and expression of the compact myocardial gene *Hey2* in published wild-type cardiac single cell transcriptomics between E10.5 and E14.5 clustered as annotated^46^. N cells: (Atrial CMs = 1340, Ventricular CMs = 7145, Endocardium = 497, Vascular endocardium = 171, Epicardium = 492, Fibroblast-like. = 616). **(c)** Expression of retinoic acid receptors (*Rara/b/g*) and retinoic acid retinoic X receptors (*Rxra/b/g*) in UMAPs of subsetted ventricular cardiomyocytes from (b). UMAP, Uniform Manifold Approximation and Projection. IVS, interventricular septum; LV, left ventricle; RV, right ventricle.

**Figure S6. Related to Figure 3. Blocking of retinoic acid signalling reception by the retinoic acid receptor inverse agonist BMS493 leads to hearts with bifid apex. (a)** Brightfield images of E14.5 hearts from embryos treated with the inverse pan RAR agonist BMS493 at the indicated stages. **(b)** Cleft ratio measurements of PBS treated (red), and BMS493 treated hearts for E10.5+E11.5, E11.5+E12.5, E12.5+E13.5, E11.5->E13.5 and E10.5->E14.5. Means, standard deviations and P-values (Welch’s t-test) are shown. The dotted lines indicate the 95% confidence interval of PBS treated embryos. N: PBS = 50, E10.5+E11.5 = 13, E11.5+E12.5 = 18, E12.5+E13.5 = 43, E11.5->E13.5 = 62, E10.5->E13.5 = 13. Ao, Aorta; CM, cardiomyocytes; LV, left ventricle; PT, pulmonary trunk; RV, right ventricle;

**Figure S7. Related to Figure 4. Cell type identification and differential gene expression analysis of sub-clusters in E11.5 scRNA sequencing data. (a)** Brightfield images of E11.5 *R26^RARα403^* and *Mef2c-AHF-Cre^Tg+^*;*R26^RARα403^* hearts. The dotted white line shows micro-dissected IVS region collected for scRNA sequencing **(b)** Violin plots of markers used to annotate clusters of single cells from micro-dissected IVS of *R26^RARα403^* and *Mef2c-AHF-Cre^Tg+^*;*R26^RARα403^* hearts at E11.5. **(c)** Violin plots of markers used to annotate clusters of the subsetted myocardial cells from (B). **(d-g)** Volcano plots of DEGs between control and mutant hearts for myocardial (d), endocardial (e), epicardial (f) and mesenchymal (g) clusters. Blue indicates downregulated and red upregulated genes; darker colours denote DEGs with a p-value < 10^-50^ (dotted line), while lighter colours denote a p-value < 10^-15^. Data is available in table S1. DEG, differentially expressed gene.

**Figure S8. Related to Figure 4. Single cell transcriptomic analysis at E11.5 identifies an expansion of trabecular myocardium in the septum on downregulation of retinoic acid signal reception. (a)** Violin plots of selected downregulated compact myocardial genes (*Tnni2*, *Hey2*, *Mb*, *Tnnt1*) and upregulated trabecular genes (*Nppa*, *Mest*, *Cited1*, *Gja1*, *Nppb*) in subsetted myocardial cells. Adjusted p-values: **** = < 10^-100^, *** = 10^-50^, ** = 10^-15^, * = 10^-5^. P-values are available in table S1. **(b)** High magnification views of the left- (rows 1 and 2) and right (rows 3 and 4) ventricular walls of *Hey2* (green) and *Nppa* (magenta) RNAscope labelled hearts from E10.5 to E13.5 of control (rows 1 and 3) and mutant (rows 2 and 4) hearts. **(c,d)** Quantification of left ventricular (c) and right ventricular (d) compact myocardial wall thickness of control and mutant hearts from E10.5 to E13.5. Mutants display thinner right, but not left, ventricular walls from E12.5 compared to controls; P-values (Welch’s t-test) are shown on the figure. N [Control; Mutant]: E10.5 [3;2]; E11.5 [3;3]; E12.5 [3;3]; E13.5 [3;4]. **(e)** Frontal section view of E12.5 cleared lightsheet imaged *R26^RARα403^*;*Cx40^GFP^*and *Mef2c-AHF-Cre^Tg+^*;*R26^RARα403^*;*Cx40^GFP^* hearts. Yellow arrowheads indicate ectopic trabeculation in the mutant septum. **(f)** Antibody staining for GFP (green) and CD31 (magenta) of 50μm vibratome sections of *R26^RARα403^*;*Cx40^GFP^* and *Mef2c-AHF-Cre^Tg+^*;*R26^RARα403^*;*Cx40^GFP^* E18.5 hearts from Figure 5B. Coronary arteries are GFP^+^; PECAM^+^ (white full arrowheads), while Cx40+ trabecular cells are GFP^+^; PECAM^-^ (empty arrowheads). Ctrl, Control; DEG, differentially expressed gene; LV, left ventricle; Mut, Mutant; RV, right ventricle.

**Figure S9. Related to Figure 5. Expanded trabecular myocardium in the forming septum on downregulation of retinoic acid signal reception. (a)**. Frontal section of nuclear and membrane labelled *R26^RARα403/mGFP^* and *Mef2c-AHF-Cre^Tg+^*;*R26^RARα403/mGFP^* hearts imaged at E10.5, E11.5 and E12.5 by lightsheet microscopy after clearing. Note the presence of an RA-independent region (arrowheads) in the septal primordium at E10.5 and at the top of the compact septum. **(b)** 2/5 wild-type hearts do not have a completely compacted septal core at E11.5. **(c)** RNAscope ISH showing *Dcc* expression in the septal primordium of *wild-type* hearts at E10.5.

**Figure S10. Related to Figure 7. Subclustering analysis of left and right myocardial cells at E11.5. (a)** Workflow showing sub-clustering of subsetted compact myocardial cells into left- and right ventricular cells. **(b)** Feature plots of genes normally associated with left ventricle (*Tbx5*, *Tbx18*) and right ventricle (*Pitx2*, *Tnni2*). **(c,d)** Volcano plots of differentially expressed genes between control and mutants of the subsetted right ventricular (c) and left ventricular (d) clusters. Blue indicates downregulated and red upregulated genes; darker colours denote p-values < 10^-20^, while lighter colours denote p-values < 10^-5^ (dotted line). Data is available in Table S1. **(e)** Violin plots of cardiac maturation- and disease related DEGs that are downregulated in mutant right ventricular compact myocardial cells. P-values are available in Table S1. **(f)** Violin plots of signalling related DEGs showing expression changes in mutant right ventricular compact myocardial cells. P-values are available in Table S1. **(g)** Wholemount RNAscope ISH for *Sfrp1* (white) followed by cleared lightsheet imaging of E11.5 *R26^RARα403^* and *Mef2c-AHF-Cre^Tg+^*;*R26^RARα403^* hearts. Full arrowheads (yellow) points to expression in the right ventricle of the control, while empty arrowheads in mutants indicate loss of expression. **(h,i)** Quantifications of signal ratio between right- and left ventricle and IVS and left ventricle for *Lpar3* (h) and *Sfrp1* (i) from images shown in Figure 7d,e. P-values (Welch’s t-test) are shown on the figures. N (all conditions) : [4 controls; 4 mutants]. CM, compact myocardium; DEG, differentially expressed genes; UMAP, Uniform Manifold Approximation and Projection.

**Figure S11. Related to Figure 8. Evaluation of vascular and differentiation defects in fetal hearts on downregulation of retinoic acid signal reception in myocardium. (a)** Brightfield images of E13.5 *R26^RARα403^* and *Mef2c-AHF-Cre^Tg+^*; *R26^RARα403^* hearts. The dotted white line shows the micro-dissected IVS regions collected for bulk RNA sequencing and RT-qPCR. **(b)** Normalized read counts of *Cre* transcript as well as genes representing different categories identified through the GSEA. P-values (adjusted p-values, DEseq2, n = [3 controls; 3 mutants]) are shown on the figure. **(c)** Brightfield images of E12.5 and E14.5 *R26^RARα403^* and *Mef2c-AHF-Cre^Tg+^*; *R26^RARα403^* hearts. Dotted white lines show micro-dissected IVS regions collected for RT-qPCR. (d, e) *Nrg1* RNAscope labelling in frontal sections of E11.5 (d) and E13.5 (e) hearts in control *R26^RARα403^* and mutant *Mef2c-AHF-Cre^Tg+^*; *R26^RARα403^* hearts. White arrowhead indicates a zone of increased expression observed in mutants, associated with the region of increased trabeculation. **(f)** Frontal section view of cleared 3D wholemount lightsheet imaged E17.5 hearts of control *R26^RARα403^*; *Cx40^GFP^*) and mutant (*Mef2c-AHF-Cre^Tg+^*; *R26^RARα403^*; *Cx40^GFP^*) samples. Empty arrowheads indicate coronary artery defects in mutants. **(g)** Frontal section of wholemount fluorescent lightsheet image of E18.5 cleared lineage traced *Mef2c-AHF-Cre^Tg+^*; *R26^mTmG^* heart showing that the septal coronary artery is not labelled by *Mef2c-AHF-Cre^Tg+^*activated GFP. IVS, interventricular septum; LV, left ventricle; RV, right ventricle; S CA, septal coronary artery.

## Supplementary Information

**Supplementary Video 1. Related to Figure 1. Lineage tracing using the *Mef2c-AHF-Cre* transgene.** 3D lightsheet imaging of cleared *Mef2c-AHF-Cre^Tg+^;R26^mTmG/+^* hearts at E11.5, E12.5 and E14.5.

**Supplementary Video 2. Related to Figure 1. 3D imaging of control and mutant hearts used for 2D measurements.** Lightsheet imaging of cleared control *R26^RARα403/+^*and mutant *Mef2c-AHF-Cre^Tg+^;R26^RARα403/+^* hearts at E14.5.

**Supplementary Video 3. Related to Figure 2. Tracing of retinoic acid responding cells in E12.5 hearts.** 3D lightsheet imaging of *RARE-CreERT2^Tg+^*;*R26^tdTomato^* hearts at E12.5 after tamoxifen injection at E10.5. tdTomato^+^ cells are observed in the ventricular septum.

**Supplementary Video 4. Related to Figure 5. Ectopic trabeculation is observed in mutant hearts from E11.5.** 3D lightsheet imaging of RNAscope labelled hearts showing the expression of a trabecular gene, *Nppa* (magenta), and a compact myocardial gene, *Hey2* (green) at E10.5 and E11.5 in control *R26^RARα403/+^* and mutant *Mef2c-AHF-Cre^Tg+^;R26^RARα403/+^*. At E10.5 the control and mutant hearts are similar, but from E11.5 there is an increased trabeculae labelling in the mutant septum.

**Supplementary Video 5. Related to Figure S9. Formation of a compact interventricular septal core at E11.5 fails in mutants.** 3D lightsheet imaging of two control and one mutant hearts at E11.5 expressing a constitutive membrane GFP marker. Most control hearts have well defined compact septum at E11.5, however some early hearts will display extended trabeculae in the basal part of the septum. All mutant hearts have ectopic trabeculae.

**Supplementary Video 6. Related to Figure 6. The septal apex is marked by *Dcc* at E11.5 and is not affected in mutants.** 3D lightsheet imaging of RNAscope labelled control (*R26^RARα403/+^*) and mutant (*Mef2c-AHF-Cre^Tg+^;R26^RARα403/+^*) E11.5 hearts showing the distribution of transcripts for the Netrin receptor *Dcc*.

**Supplementary Video 7. Related to Figure S10. *Sfrp1* expression is downregulated in the septum and right ventricle of mutant hearts.** 3D wholemount light sheet images of control (*R26^RARα403/+^*) and mutant (*Mef2c-AHF-Cre^Tg+^;R26^RARα403/+^*) E11.5 hearts after RNAscope hybridisation for *Sfrp1* expression, showing downregulation in the right ventricle and septum, but not the left ventricle, of mutant hearts.

**Table S1. List of differentially expressed genes between control and mutant cell clusters from microdissected E11.5 IVS regions of *R26^RARα403^* and *Mef2c-AHF-Cre^Tg+^*;*R26^RARα403^* hearts.**

**Table S2. List of top 100 positive markers for the *Dcc^high^* inlet septum myocardial cell cluster.**

**Table S3. List of differentially expressed genes calculated by DEseq2 analysis of bulk RNA sequencing of E13.5 IVS of control *R26^RARα403^* and mutant *Mef2c-AHF-Cre^Tg+^*;*R26^RARα403^* hearts.**

**Table S4. List of GO terms from the gene set enrichment analysis of E13.5 IVS of control *R26^RARα403^* and mutant *Mef2c-AHF-Cre^Tg+^*;*R26^RARα403^* hearts.**

**Table S5. List of primers used for genotyping and RT-qPCR.**

