## Supplementary material for "Morphogenesis of the muscular ventricular septum of the mouse heart is driven by retinoic acid signalling to myocardium": Bonnelykke et al Supplementary Figures

Figure S1 - Related to Figure 1 Bønnelykke *et al.*

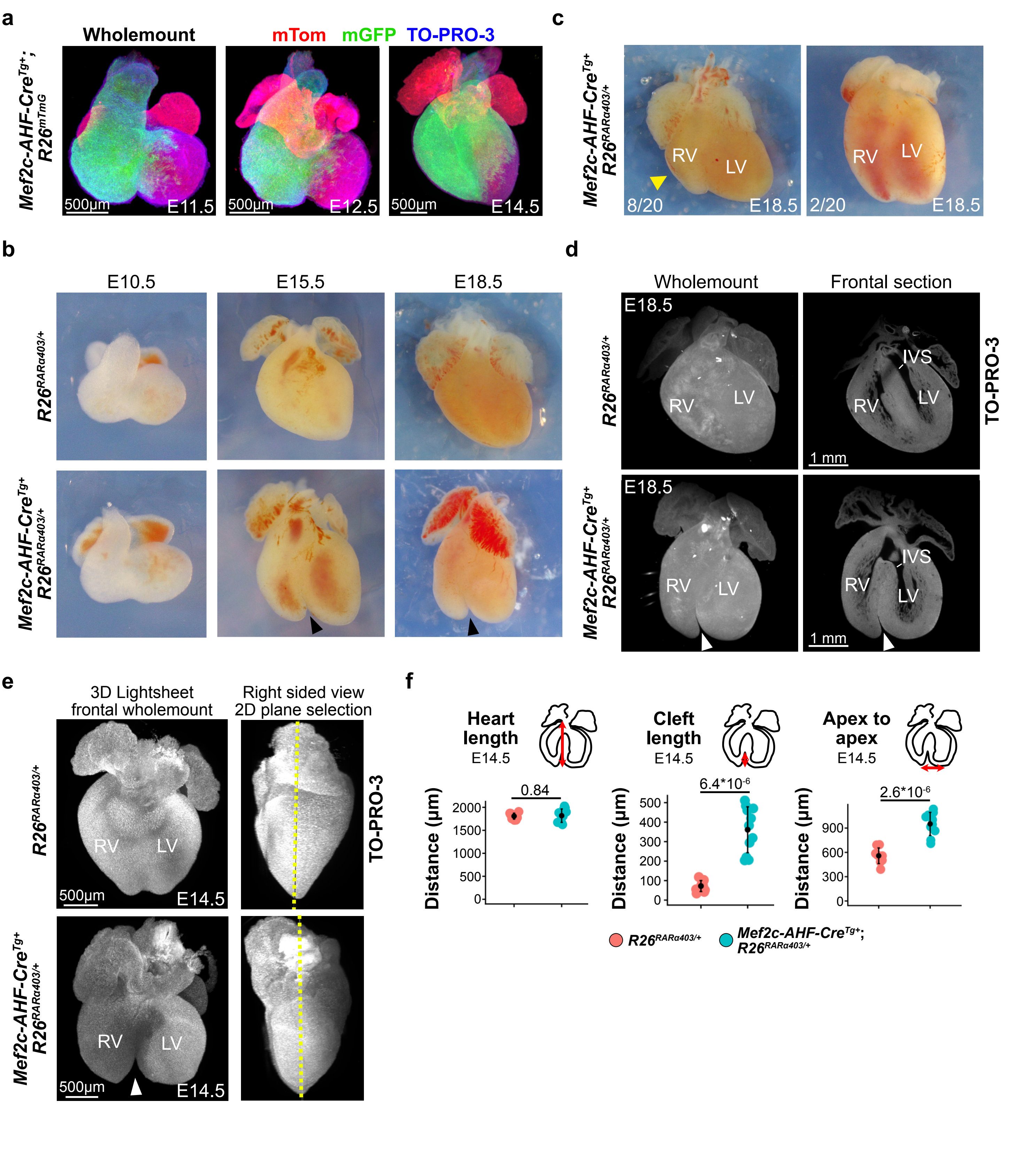

**a**

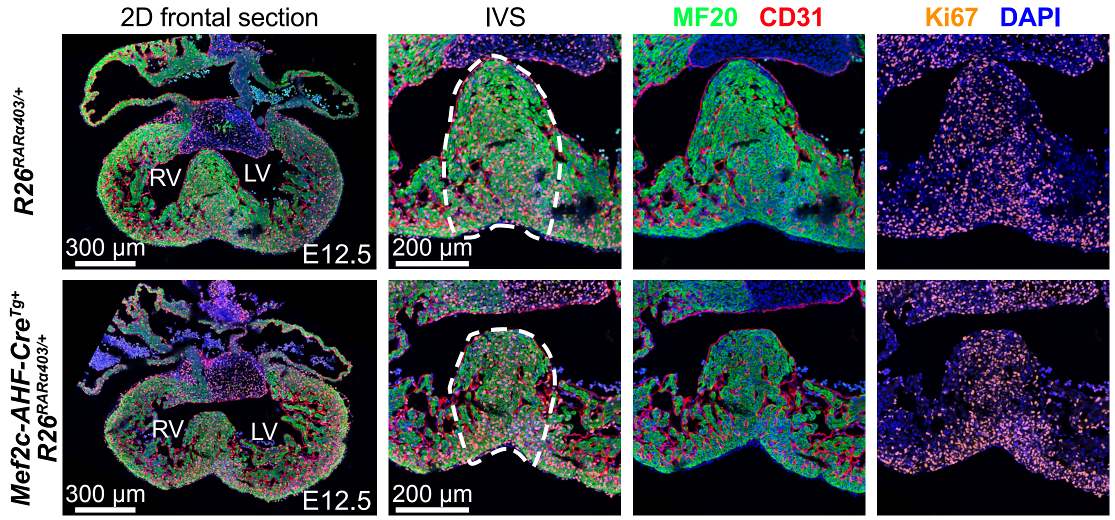
2D frontal section
IVS
MF20 CD31
Ki67 DAPI
R26RARα403/+
Mef2c-AHF-CreTg+ R26RARα403/+
RV
LV
300 µm
200 µm
E12.5

**b**

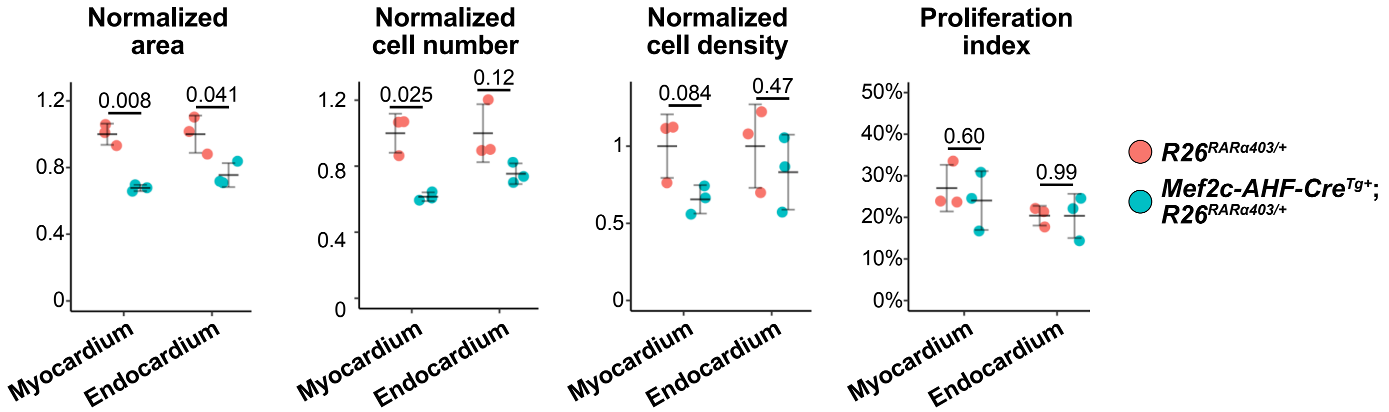
Normalized area
Normalized cell number
Normalized cell density
Proliferation index
0.008
0.041
0.025
0.12
0.084
0.47
0.60
0.99
1.2
0.8
0.4
0
1
0.5
50%
40%
30%
20%
10%
0%
Myocardium
Endocardium
R26RARα403/+
Mef2c-AHF-CreTg+; R26RARα403/+

**c**

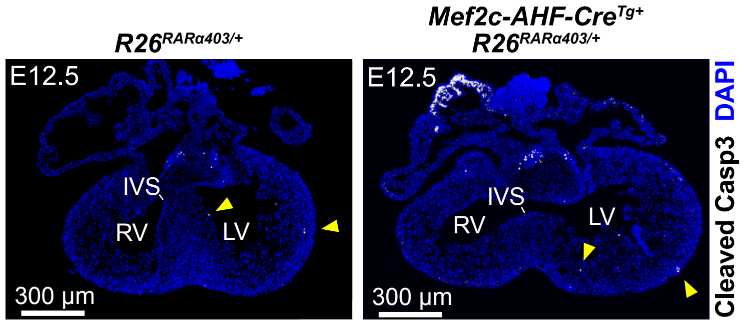
R26RARα403/+
Mef2c-AHF-CreTg+ R26RARα403/+
E12.5
IVS
RV
LV
300 µm
Cleaved Casp3 DAPI

### Figure S3 - Related to Figure 1

Bønnelykke *et al.*

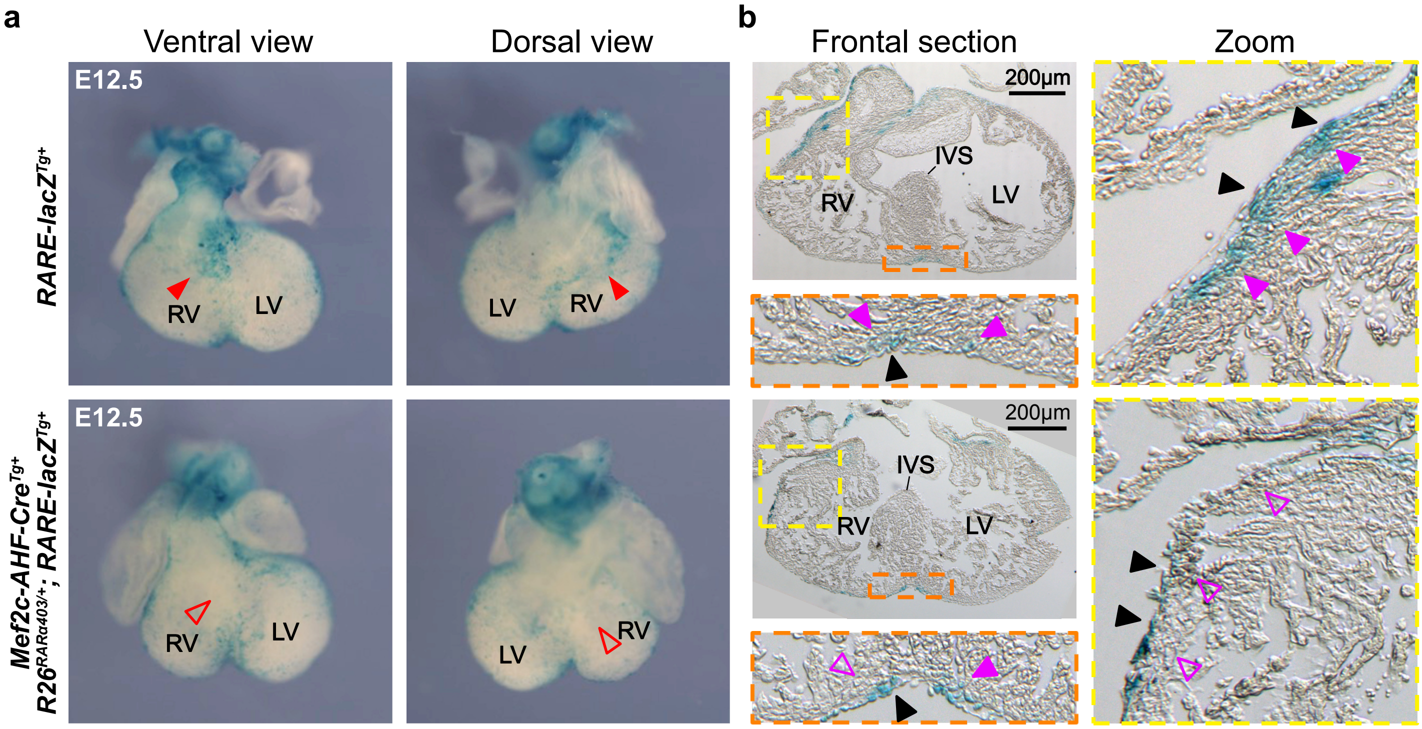

Figure S4 - Related to Figure 2 Bønnelykke *et al.*

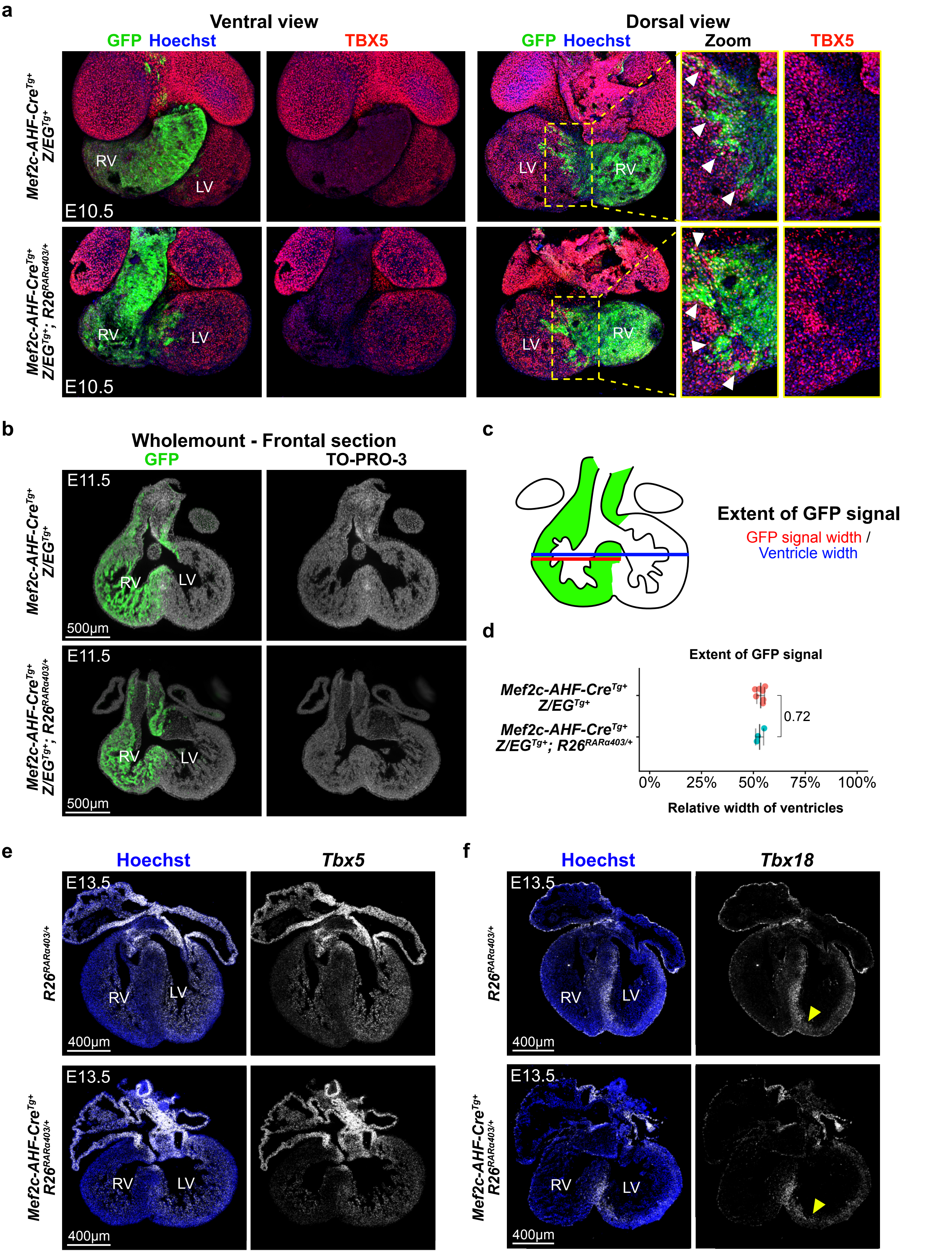

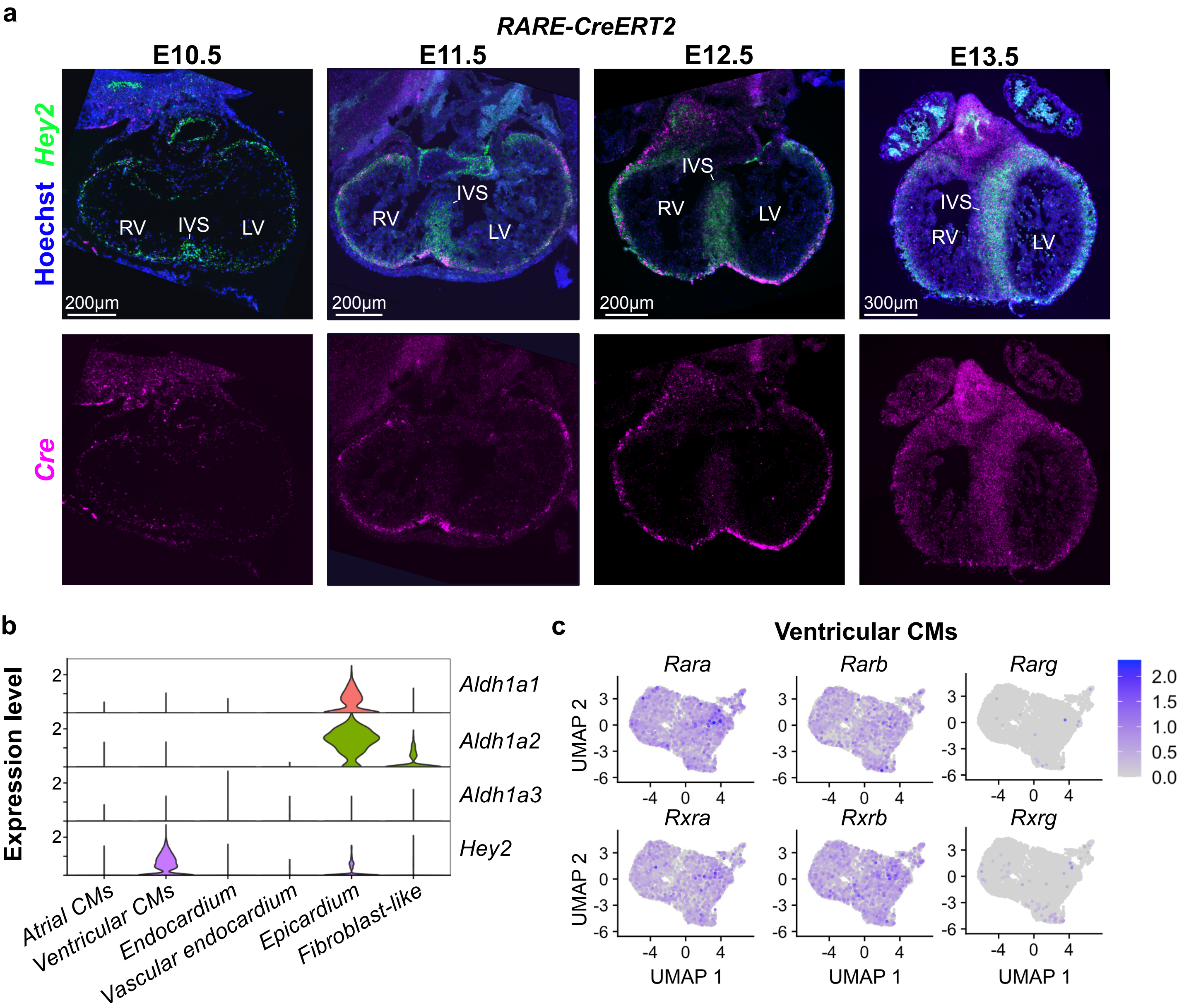
a
RARE-CreERT2
E10.5
E11.5
E12.5
E13.5
Hoechst Hey2
Cre
RV
IVS
LV
200µm
300µm
b
Expression level
Aldh1a1
Aldh1a2
Aldh1a3
Hey2
Atrial CMs
Ventricular CMs
Endocardium
Vascular endocardium
Epicardium
Fibroblast-like
c
Ventricular CMs
Rara
Rarb
Rarg
Rxra
Rxrb
Rxrg
UMAP 2
UMAP 1
2.0
1.5
1.0
0.5
0.0

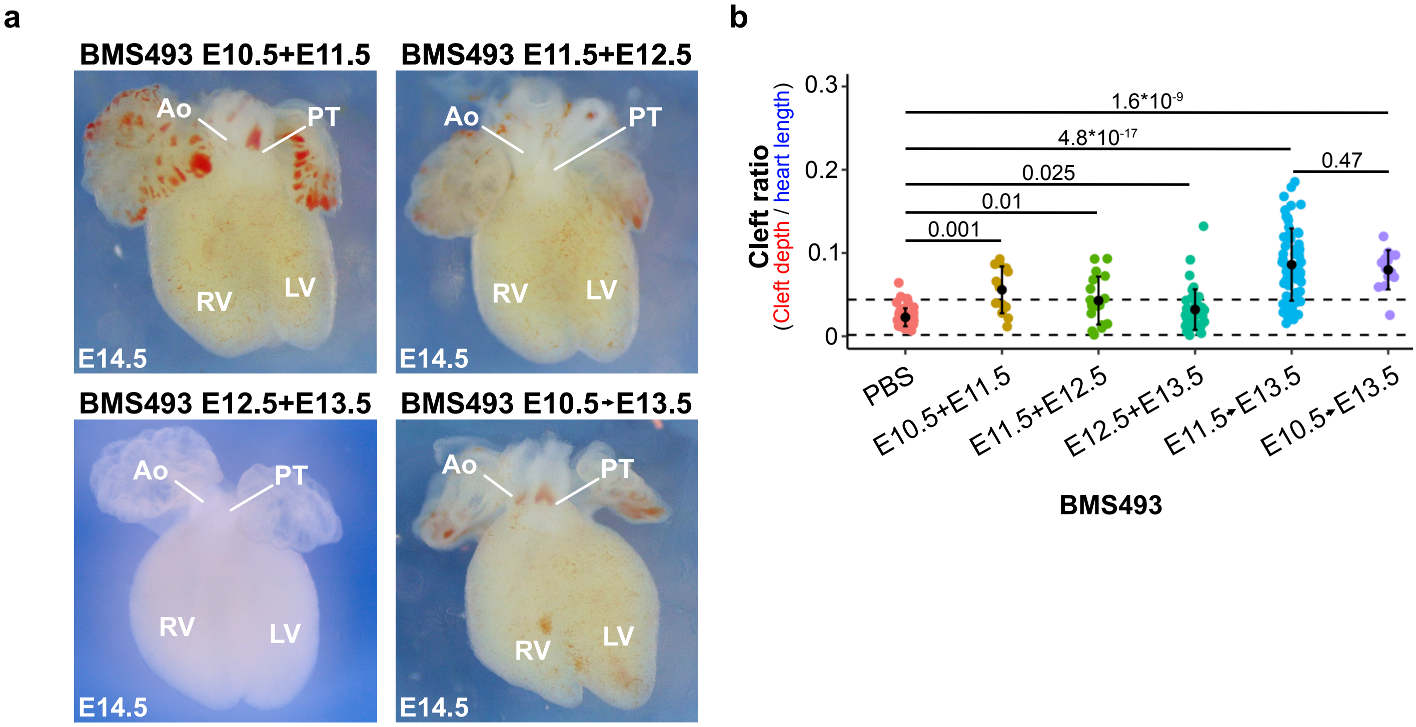
a
BMS493 E10.5+E11.5
BMS493 E11.5+E12.5
BMS493 E12.5+E13.5
BMS493 E10.5→E13.5
Ao
PT
RV
LV
E14.5
b
Cleft ratio
(Cleft depth / heart length)
1.6*10-9
4.8*10-17
0.025
0.01
0.001
0.47
0.3
0.2
0.1
0
PBS
E10.5+E11.5
E11.5+E12.5
E12.5+E13.5
E11.5→E13.5
E10.5→E13.5
BMS493

### Figure S7 - Related to Figure 4

Bønnelykke *et al.*

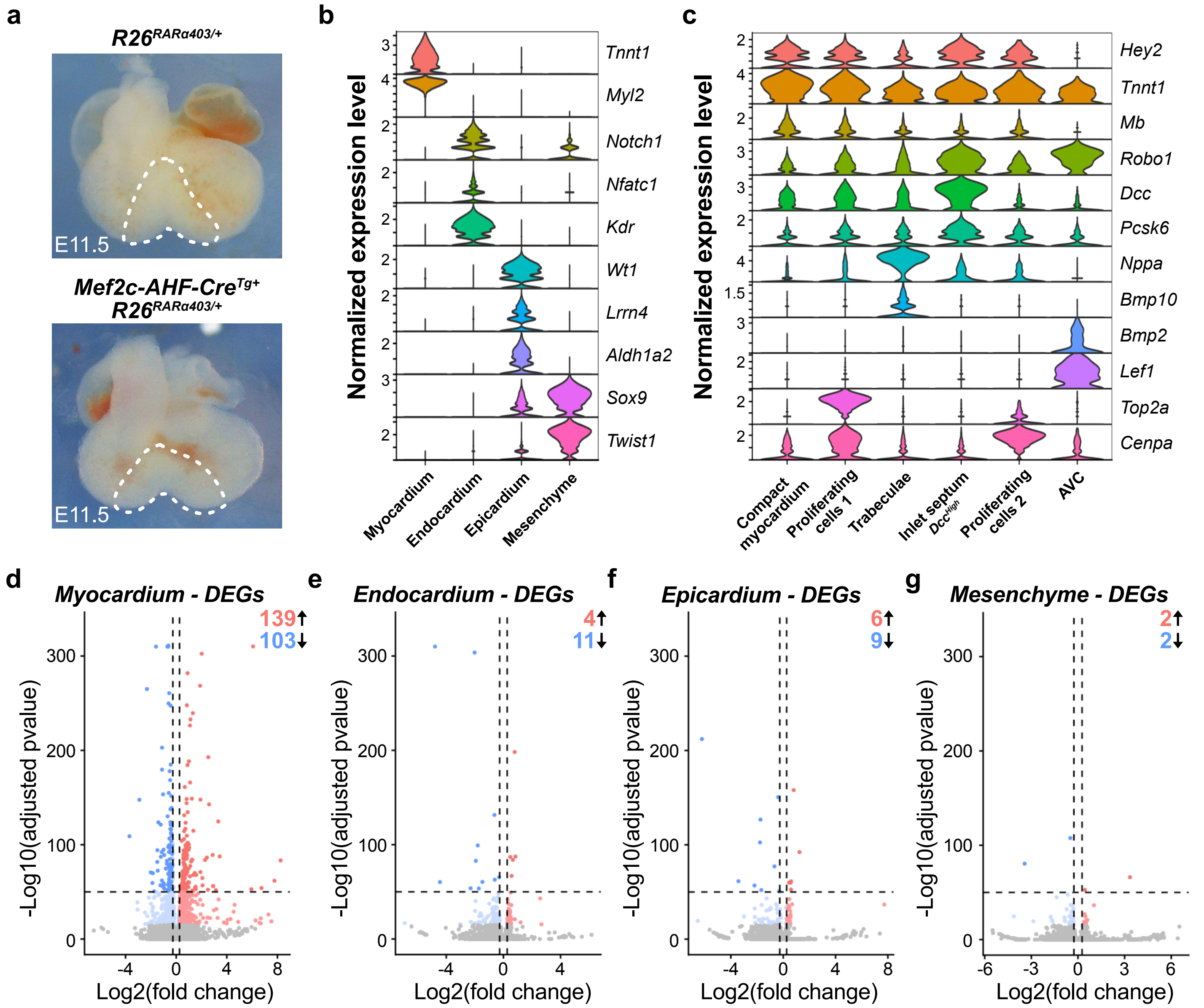

Figure S8 - Related to Figure 4 and Figure 5 Bønnelykke *et al.*

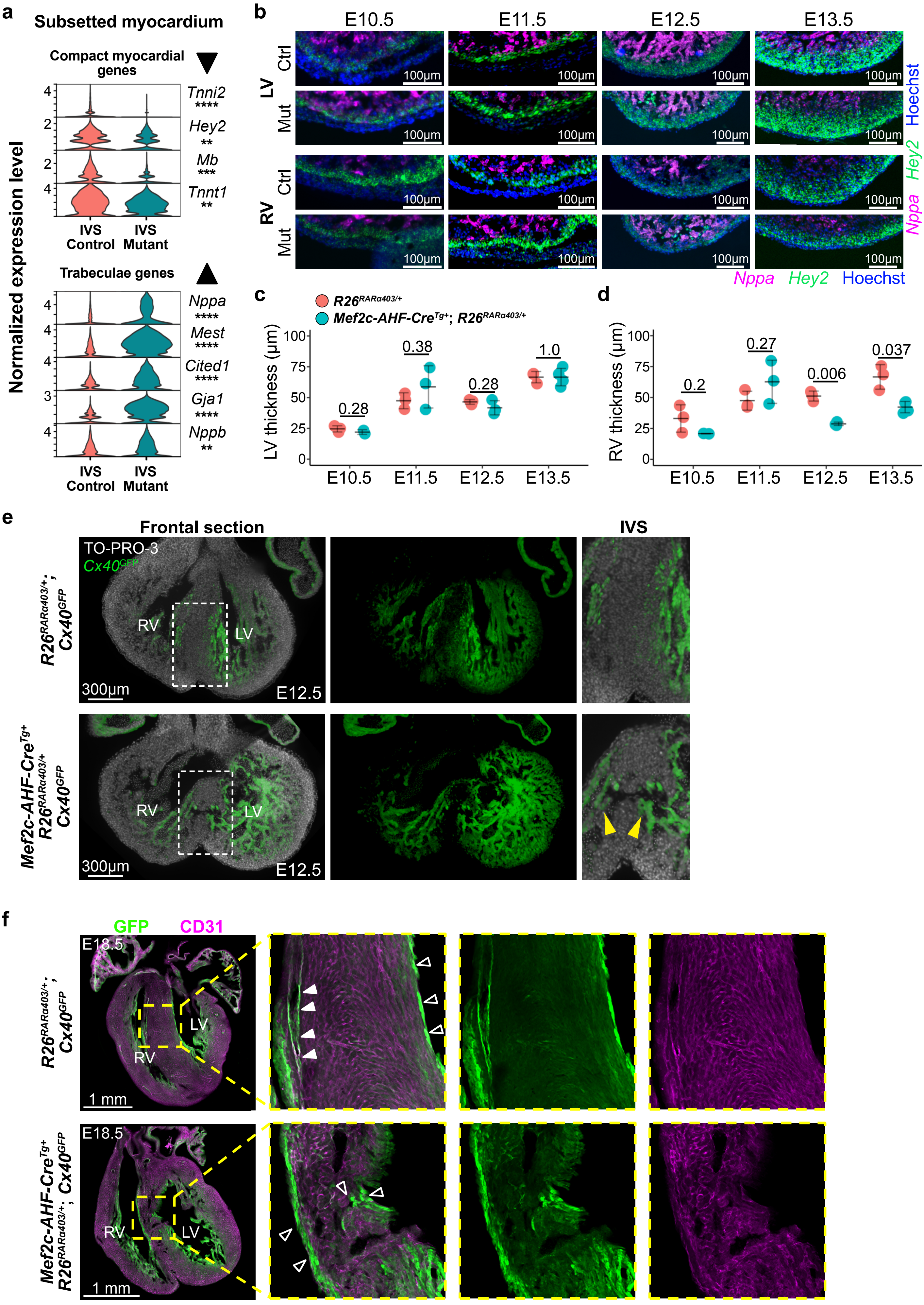

### Figure S9 - Related to Figure 5 and Figure 6



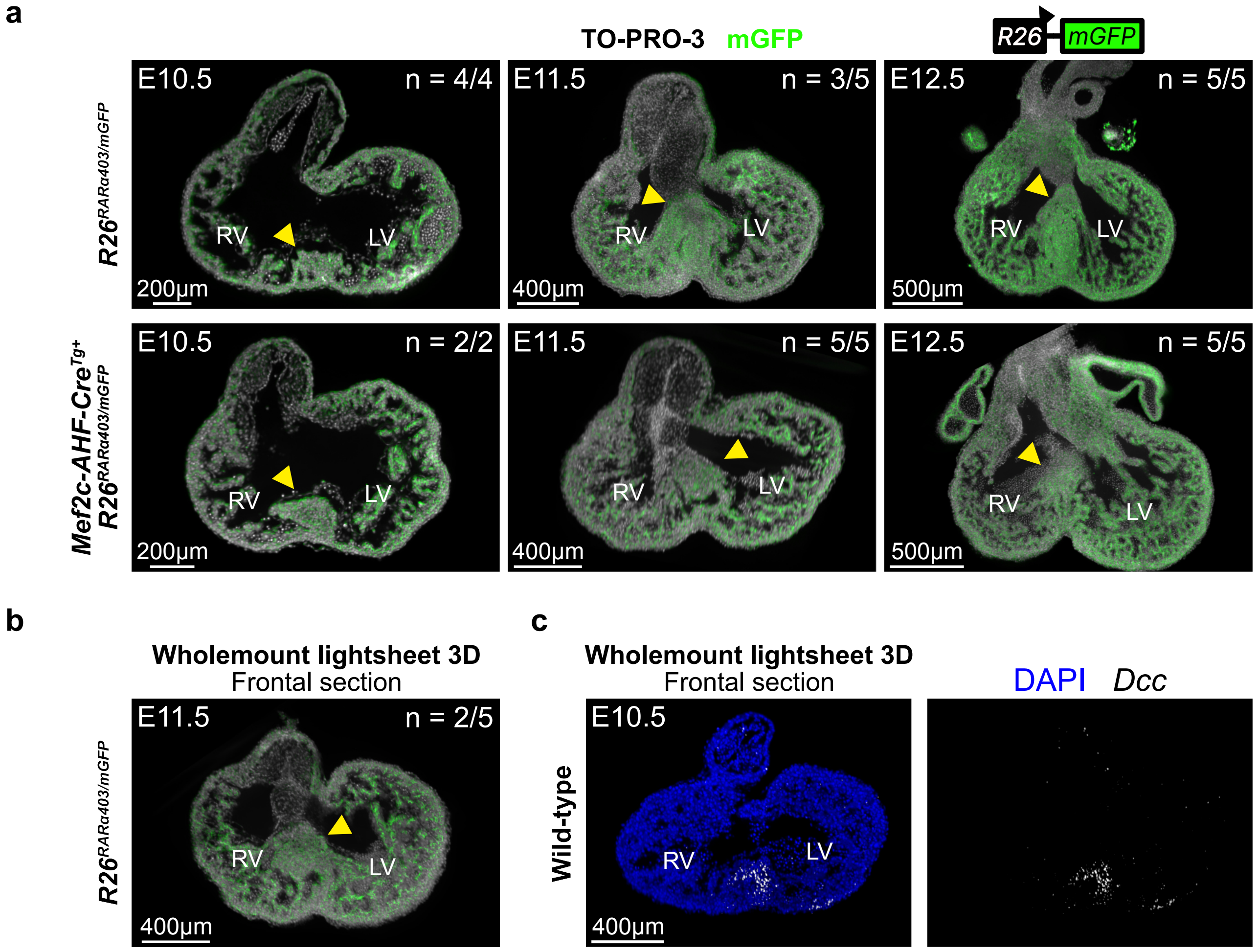

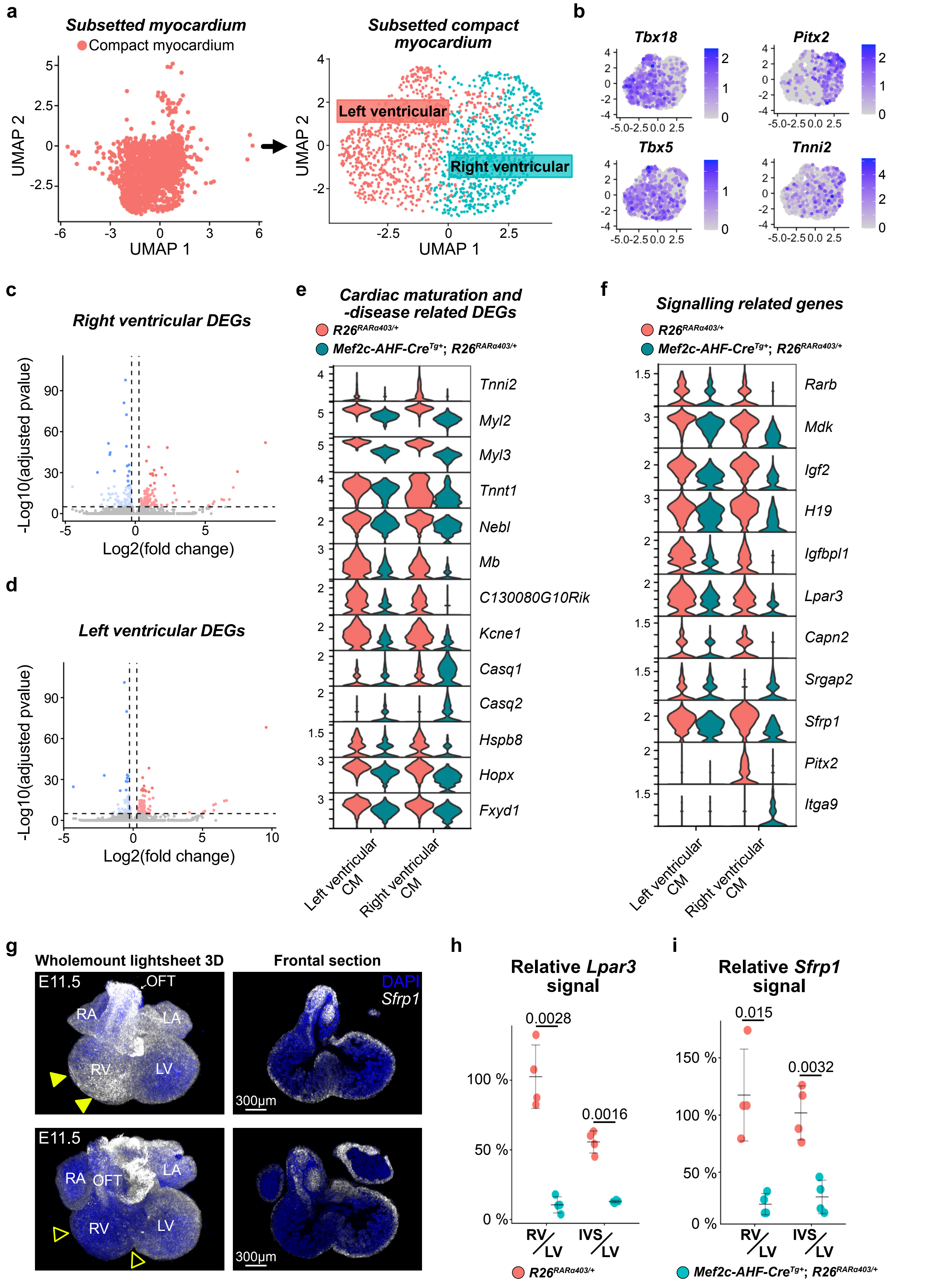
a
Subsetted myocardium
Compact myocardium
UMAP 1
UMAP 2
Subsetted compact myocardium
Left ventricular
Right ventricular
b
Tbx18
Pitx2
Tbx5
Tnni2
c
Right ventricular DEGs
-Log10(adjusted pvalue)
Log2(fold change)
d
Left ventricular DEGs
e
Cardiac maturation and -disease related DEGs
R26RARα403/+
Mef2c-AHF-CreTg+; R26RARα403/+
Tnni2
Myl2
Myl3
Tnnt1
Nebl
Mb
C130080G10Rik
Kcne1
Casq1
Casq2
Hspb8
Hopx
Fxyd1
Left ventricular CM
Right ventricular CM
f
Signalling related genes
Rarb
Mdk
Igf2
H19
Igfbpl1
Lpar3
Capn2
Srgap2
Sfrp1
Pitx2
Itga9
g
Wholemount lightsheet 3D
Frontal section
E11.5
OFT
RA
LA
RV
LV
DAPI
Sfrp1
300µm
h
Relative Lpar3 signal
0.0028
0.0016
RV/LV
IVS/LV
i
Relative Sfrp1 signal
0.015
0.0032

### Figure S11 - Related to Figure 8

Bønnelykke *et al.*

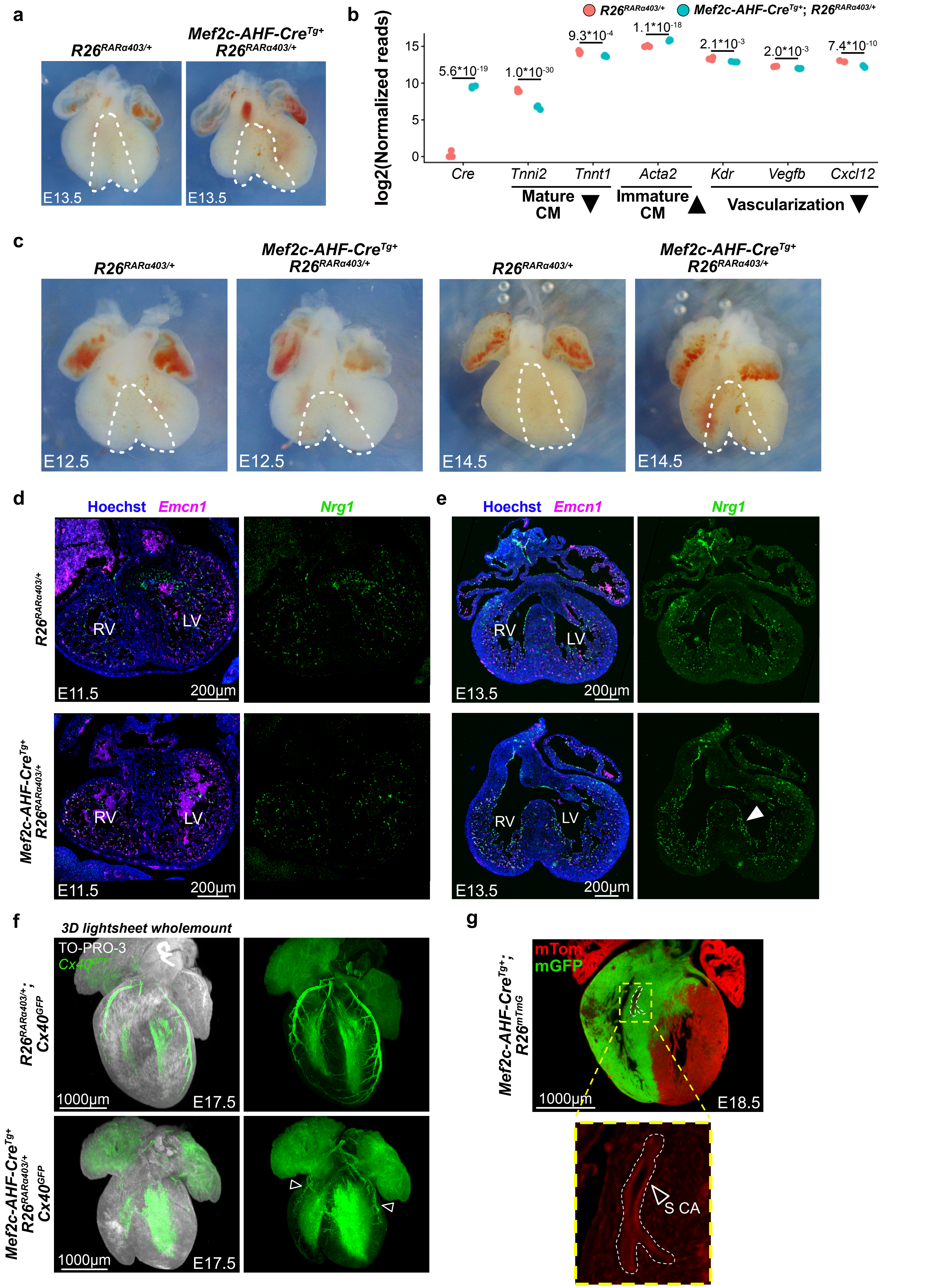
